# Optimising the flow of mechanical energy in musculoskeletal systems through gearing

**DOI:** 10.1101/2024.04.05.588347

**Authors:** D T Polet, D Labonte

## Abstract

Movement is integral to animal life, and most animal movement is actuated by the same engine: skeletal muscle. Muscle input is typically mediated by skeletal elements, resulting in musculoskeletal systems that are “geared”: at any instant, the muscle force and velocity are related to the output force and velocity only via a proportionality constant *G*, the “mechanical advantage”. The functional analysis of such “simple machines” has traditionally centred around this instantaneous interpretation, such that a small vs large *G* is thought to reflect a fast vs forceful system, respectively. But evidence is mounting that a complete analysis ought to also consider the mechanical energy output of a complete contraction. Here, we approach this task systematically, and use the theory of physiological similarity to study how gearing affects the flow of mechanical energy in a minimalist model of a musculoskeletal system. Gearing influences the flow of mechanical energy in two key ways: it can curtail muscle work output, because it determines the ratio between the characteristic muscle work and kinetic energy capacity; and it defines how each unit of muscle work is partitioned into different system energies, i. e. into kinetic vs. “parasitic” energy such as heat. As a consequence of both effects, delivering maximum work in minimum time and with maximum transmission efficiency generally requires a mechanical advantage of intermediate magnitude. This optimality condition can be expressed in terms of two dimensionless numbers, which reflect the key geometric, physiological, and physical properties of the interrogated musculoskeletal system, and the environment in which the contraction takes place. Illustrative application to exemplar musculoskeletal systems predicts plausible mechanical advantages in disparate biomechanical scenarios; yields a speculative explanation for why gearing is typically used to attenuate the instantaneous force output (*G*_opt_ *<* 1); and predicts how *G* needs to vary systematically with animal size to optimise the delivery of mechanical energy, in superficial agreement with empirical observations. A many-to-one-mapping from musculoskeletal geometry to mechanical performance is identified, such that differences in *G* alone do not provide a reliable indicator for specialisation for force vs speed—neither instantaneously, nor in terms of mechanical energy output. The energy framework presented here can be used to estimate an optimal mechanical advantage across variable muscle physiology, anatomy, mechanical environment and animal size, and so facilitates investigation of the extent to which selection has made efficient use of gearing as degree of freedom in musculoskeletal “design”.

## Introduction

*Give me a place to stand and with a lever I will move the whole world*.

Archimedes of Syracuse

Archimedes referred to them as “simple machines”, yet their influence on human culture, history, and technology has been anything but: cycling up a hill to get to work, building a pyramid to worship a supposed deity, or opening a cold drink on a hot summer’s day—many onerous tasks are substantially eased if not enabled by contrivances that provide a “mechanical advantage”, i.e. that allow the movement of a large payload through application of a small input load. Machines that *leverage* a mechanical advantage are not just everywhere around us, but they are literally within us, for evolution has selected an engine that drives rotational movements by shortening: skeletal muscle.

Muscle is the primary animal motor, and it therefore stands to reason that understanding the physical limits to animal movement must involve an analysis of the physical and physiological constraints on muscle performance (Alexander 2003; Biewener and Patek 2018). As with Archimedes’s simple machines, a muscle’s force, shortening speed and displacement are not, in general, equal to the force, speed or displacement experienced by the object it actuates. Instead, muscle action is transmitted through joints, so that instantaneous muscle input and skeletal output are proportional to one another instead; musculoskeletal systems are generally “geared”. In modulating the effect of muscle input on skeletal output, gearing provides a “degree of freedom” on which evolutionary selection can act. But what, if any, is the best way to gear?

Musculoskeletal gearing manifests in many different forms: joints that connect skeletal segments (Borelli 1680; Gray 1944; Gregory 1912a; Smith and Savage 1956); complex kinetic linkages formed of multiple rigid elements (Anderson and Patek 2015; Hulsey and Wainwright 2002; Muñoz et al. 2017; Olsen and Westneat 2016; Westneat 1994); muscle fiber pennation (Benninghoff and Rollhäuser 1952; Brainerd and Azizi 2005; Gans and Bock 1965; Pfuhl 1937); or, indeed, interlocking elements that look unmistakably like engineered gears (Burrows and Sutton 2013). Irrespective of the implementation of gearing, its effect can be described through introduction of a “mechanical advantage”, *G*, usually defined by the ratio of two moment or lever arms, *L*. For a simple hinge joint (Figure 1), it is convenient to define these lever arms as the perpendicular distances between the joint axis of rotation and the lines of action of input and output force vectors (*L*_*i*_ and *L*_*o*_, respectively; Figure 2a), so that the rotational dynamics can be written as a scalar expression. The proportionality constants that link instantaneous output (*F*_*o*_) and input force (*F*_*i*_), instantaneous input (*v*) and output velocity (*u*), and the input (*δ*) and output displacement (*x*) are then equal to the ratio between an effective “in-lever” (*L*_*i*_), and an effective “out-lever” (*L*_*o*_):

**Figure 1:**
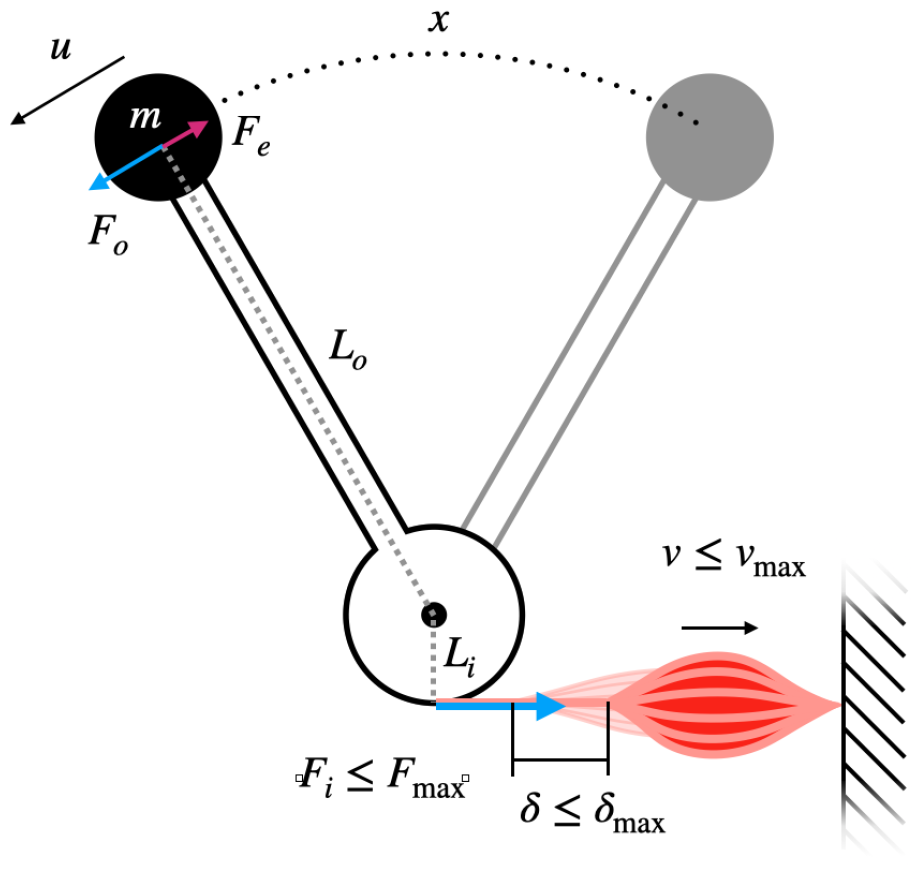
Schematic of the idealised musculoskeletal system that it studied in this paper. The muscle operates with constant mechanical advantage *G* = *L*_*i*_*/L*_*o*_, has a maximum contraction velocity *v*_max_, and a maximum displacement capacity *δ*_max_. Once either limit is exceeded, muscle tensile force instantaneously drops from its maximum value *F*_*i*_(*v ≤ v*_max_, *δ ≤ δ*_max_) = *F*_max_ to zero, *F*_*i*_(*v > v*_max_, *δ > δ*_max_) = 0. The inertia associated with the mass of the lever is assumed small relative to the inertia associated with the payload of mass *m*. External forces *F*_*e*_ may resist motion, such that *F*_net_ = *F*_*i*_ *− F*_*e*_.

**Figure 2:**
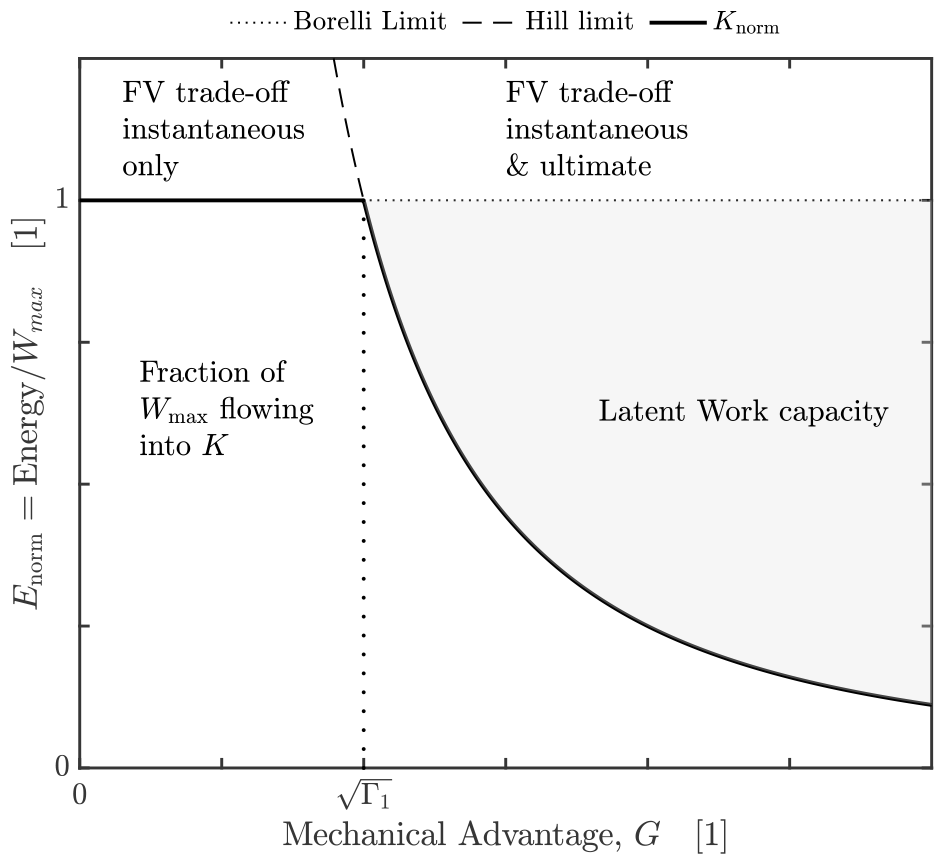
The energy landscape for an idealised musculoskeletal system in an inertial contraction, *F*_net_ *≈ F*_*o*_. Because all external forces are negligible, all muscle work is converted into kinetic energy, *W*_*m*_ = *K*. For a sufficiently small mechanical advantage, 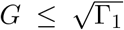, the muscle can deliver its maximum work capacity, defined by the Borellilimit *W*_*m*_ = *W*_max_. In this regime, variations in *G* merely change the time over which *W*_max_ is delivered; force-velocity trade-offs only apply instantaneously, but do not map on the outcome of the contraction (Labonte 2023; McHenry 2010). For 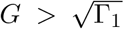, however, the muscle contraction speed reaches *v*_max_ before the muscle has delivered *W*_max_, so that muscle energy output is bound by the Hill-limit instead, *W*_*m*_ = *K*_max_ *< W*_max_. A fraction 1 *−* Γ of its work capacity remains “latent” in this regime, in the sense that the system can access it principle, for example, if the payload is increased (which changes Γ_1_). In this regime, force-velocity trade-offs apply instantaneously, and map onto the ultimate outcome of the contraction Arnold et al. (2011); Labonte (2023). 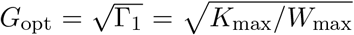 thus represents the optimal mechanical advantage in an inertial contraction: the muscle can deliver maximum kinetic energy in minimum time. The separation of the energy landscapes into these two regimes reconciles the debate between McHenry, Arnold and others Arnold et al. (2011); McHenry and Summers (2011); McHenry (2010, 2012): both arguments are correct, but in opposite regimes of the physiological similarity index Γ = Γ_1_*G*^*−*2^ (see text for details).

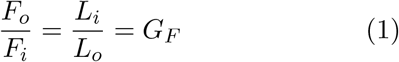

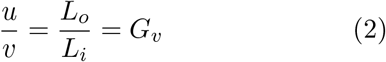

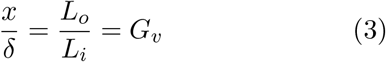

where we made the common assumption that the lever itself is of negligible mass, and note that the same musculoskeletal systems can operate with different combinations of the “moment arms” *L*_*i*_ and *L*_*o*_, for example by altering the position of food along a jaw and the jaw opening angle. The velocity (or displacement) advantage, 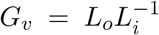 is used by engineers to parameterise gears and appears occasionally in the biomechanical literature (e.g. Azizi and Brainerd 2007; Carrier et al. 1998a; McHenry 2012); the force advantage, 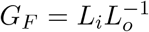, in turn, is more common in the biomechanical literature, perhaps because of its historical origin in the work of Borelli (Borelli 1680). For convenience, one of the two ratios is typically referred to as mechanical advantage or gear ratio, so that the other follows as the inverse; throughout this text, we will define the mechanical advantage as the force advantage, in keeping with much of the biomechanical literature, 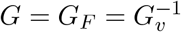.

Three properties of eq. 1 have catapulted *G* into arguably the dominant functional parameter in the study of skeletal functional morphology. First, it is mathematically simple. Second, it appears to permit comparative inference about musculoskeletal dynamics, a complex trait, through measurement of two anatomical traits that are often readily accessible. And third, despite this simplicity, the implied functional significance appears both straightforward and significant: *G* is the force amplification factor, and the inverse of the velocity amplification factor— eqs. 1-2 encode a trade-off between amplifying force or velocity. As Galileo put it, “what is lost in force is gained in speed”. The magnitude of *G* may consequently be interpreted as an indication for how this trade-off has been resolved, i. e. whether selection has acted to increase output force or speed. Because trade-offs can act both as constraints and drivers of evolutionary diversification (Arnold 1992; Burress and Muñoz 2023; Garland et al. 2022; Herrel et al. 2009; Holzman et al. 2011), *G* has played a key role in a rich body of literature on the macroevolutionary diversification of skeletal anatomy.

Early studies of skeletal anatomy considered the mechanical function encoded by *G* as an essential evolutionary driver of postural variations in ungulates (Gregory 1912b); as determinant of limb function during locomotion (Gray 1944); and as primary explanation for the diversity of mammalian skeletal shape in general (Smith and Savage 1956). In the last three decades alone, *G* has been the principal functional metric in the study of the ecological speciation in Darwin’s finches (Herrel et al. 2009); the evolution of feeding mechanisms in fish (Westneat 1994, 2004); the evolution of skull form in bats (Dumont et al. 2014; Santana et al. 2012); the morphological diversification of jaw morphology in vertebrates (Anderson et al. 2011; Deakin et al. 2023) and of chela morphology in scorpions (Simone and van Der Meijden 2018); bite force estimation in extinct species (Sakamoto et al. 2010, 2019); the early evolution of biting-chewing performance in the hexapoda (Blanke 2019); the ecological and taxonomic diversification of crown mammals (Tseng et al. 2023), and the diet of Mesozoic mammals (Morales-García et al. 2021); and the variability of rates of phenoptypic evolution in cichlids (Burress and Muñoz 2023). There is probably no other biomechanical metric that has been measured as extensively, and that has served as foundation for such far reaching comparative, ecological and evolutionary conclusions.

### Is the force-velocity trade-off universal? A shift in perspective

The functional interpretation of *G* as arbiter of a force-velocity trade-off continues to dominate the literature, but it is neither the only interpretation, nor is it universally accepted. Alternative but not mutually exclusive suggestions posit that the magnitude of *G* influences the efficiency of locomotion (Reilly et al. 2007; Ren et al. 2010; Usherwood 2013); that systematic variations in *G* with animal body size help ensure that static equilibrium can be maintained with equal muscle effort despite the systematic variation of the muscle to weight force ratio (Biewener 1989a; Dick and Clemente 2017); or that an appropriate choice and dynamic variation of *G* can optimise muscle mechanical performance during running (Carrier et al. 1998b, 1994), and maximise the mechanical energy than can be stored in elastic elements to enhance performance during explosive movements (Olberding et al. 2019; Roberts and Marsh 2003). Dissenting interpretations, questioning whether *G* encodes a universal force-velocity trade-off, have existed since at least the late 1970s (Coombs 1978), and have grown both louder and more numerous in the last decade (McHenry and Summers 2011; McHenry 2010, 2012; Olberding et al. 2019; Osgood et al. 2021). What is the origin of the disagreement?

To understand the primary objection, note an essential subtlety in eqs. 1-2: the force- and velocity advantage apply *instantaneously*, i. e. at any specific point in time. Consider two musculoskeletal systems, identical except that one has twice the mechanical advantage of the other. The instantaneous interpretation of *G* implies that if the muscle in both systems exerts an instantaneous force *F*_*i*_ and shortens with an instantaneous velocity *v*, then the instantaneous output force and velocity differ by factors of two as well– the system with the larger gear ratio exerts a larger net force, *F*_*o*_ = 2*GF*_*i*_, but moves with a smaller velocity, *u* = *v*(2*G*)^*−*1^ and *vice versa*; varying *G* trades instantaneous output force against instantaneous output velocity. This interpretation is certainly *instantaneously* correct (Arnold et al. 2011), and is not called into question (previous misunderstandings partially arose from the use of incorrect terminology, see below and Arnold et al. 2011; McHenry and Summers 2011; McHenry 2010; Osgood et al. 2021). But how did the muscle in both systems come to achieve the shortening velocity *v*? If the muscle starts at rest, then reaching *v* demands acceleration. Intriguingly, a larger mechanical advantage will increase the output force, *F*_*o*_ = *GF*_*i*_, and with it the acceleration, *a* = *F*_*o*_*/m*, so that speed is gained more quickly and with less displacement. It is tempting, then, to ask why a larger mechanical advantage should not make it *easier* to reach a larger velocity– the exact opposite of a force-velocity tradeoff.

The usual counterpoint to this objection nicely mirrors the historical development of the understanding of “simple machines”: for Aristotle and Archimedes, to use a lever was to defy the laws of nature through the use of art, for a large weight could be made to move with minuscule force (Gatto 2017). Heron, in his *Mechanics*, was perhaps the first Western scholar to note that the decrease in required forces comes with a cost Gatto (2017), but it was Galileo who firmly established this insight through the introduction of the concepts of power and work: instead of in “a certain manner deceiving nature”, gearing may reduce the force required to move an object, but it cannot alter the work a machine can do. It is this focus on muscle work, the integral of force with respect to displacement, that has called the universality of force-velocity trade-offs into question (McHenry and Summers 2011; McHenry 2010, 2012; Olberding et al. 2019; Osgood et al. 2021). Instead of asking “For a given instantaneous muscle force *F*_*i*_, shortening speed *v* and gear ratio *G*, what are the instantaneous output force *F*_*o*_ and velocity *u*?”, as supposed by the instantaneous perspective, an energy perspective on gearing asks: “How does gearing influence the ability of muscle to do work, and how does it control the ‘transmission efficiency’ of muscle work into the kinetic energy?” (McHenry and Summers 2011; McHenry 2010, 2012; Olberding et al. 2019; Osgood et al. 2021). In other words, instantaneous quantities, such as force and velocity, are replaced with integral quantities, such as work or impulse; the focus lies no longer on a specific instant in time during a contraction, but on the contraction outcome. The key problem is that these two perspectives can be in conflict.

This conflict came to the fore in a series of papers published in the 2010s. In what was perhaps the first study to redirect the focus from an instantaneous interpretation of gearing to one that analysed energy flow, McHenry (2010) argued that “there is no force-velocity trade-off in dynamic lever systems”, and, consequently, that “skeletal geometry provides a necessary, but insufficient, means to characterize the dynamic performance of a lever system”. Arnold et al. (2011) countered, in no less certain terms, that “there is always a trade-off between speed and force in a lever system”, and that the results of McHenry merely reflected the trivial conservation of energy. This critical counterargument was partially triggered by the use of “quasi-static” and “static equilibrium” by McHenry, although “instantaneous” was meant. McHenry and Summers (2011) clarified that the disagreement largely reflects one of emphasis and not of principle : force-velocity trade-offs exist instantaneously, as argued by Arnold et al. (2011), but they are not absolute, for they disappear when the work output of muscle is kept constant— instantaneous amplification does not necessarily map onto the outcome of a contraction (McHenry and Summers 2011). But Arnold et al. also questioned the assumption that muscle work input can be considered constant, and instead suggested that the mechanical metric of interest was muscle power, so that a force-velocity trade-off also persists dynamically (Arnold et al. 2011). McHenry then went one step further, and demonstrated more formally what was already suggested in the discussion of the initial commentary: the velocity resulting from a contraction can vary with gear ratio *even when muscle supplies a constant amount of energy*, because gearing controls the “transmission efficiency” of the energy conversion, i. e. how much of each unit of muscle work serves to increase kinetic energy (McHenry 2012, This is unpacked more formally below). Using forward dynamic simulations, McHenry showed elegantly that variations in gear ratio can leave the resulting speed unaffected, decrease it, or increase it, depending on the “mechanical environment”; there exists no one-to-one mapping between gear ratio and output speed, and variations in *G* are neither necessary nor sufficient to change the maximum speed a musculoskeletal system can impart (see also Coombs 1978; Labonte 2023; Labonte et al. 2024; Lee et al. 2009; Nagano and Komura 2003; Olberding et al. 2019; Osgood et al. 2021; Richards 2011; Richards and Clemente 2013a, for related results).

Here, we take McHenry’s work as a starting point, and seek to develop a first principles framework to analyse the influence of the mechanical advantage on the mechanical energy output and mechanical energy flow in musculoskeletal systems. Our analysis builds on the theory of physiological similarity (Labonte 2023), and differs from earlier work in two key aspects: First, instead of an “elastic actuator” akin to a spring, we will analyse the dynamics of a system driven directly by muscle. This difference will enable us to show that variations in mechanical advantage can control how much work muscle can deliver, so reconciling Arnold et al.’s objection with McHenry’s observation. Second, where others relied on numerical simulations (McHenry 2010, 2012; Olberding et al. 2019; Osgood et al. 2021), we will derive exact symbolic relationships wherever possible. By leaning on explicit analytical expressions rather than simulations, we will be able to derive results that readily generalise, and sharpen our understanding of the physical principles at play. The aims of this work are to unambiguously pinpoint the role of gearing in modulating muscle energy output through a rigorous mechanical analysis; to investigate how gearing interacts with external forces to modulate the conversion of muscle work input into kinetic energy output; and to further clarify the conditions under which variations in *G* can be interpreted as an evolutionary adaptation that reflects force-velocity trade-offs across animal size and mechanical environment.

## Model formulation and assumptions

The aim of this manuscript is to analyse how gearing influences (i) the work muscle can deliver in a single contraction, and (ii) how it controls the distribution of each unit of muscle work among different forms of energy that characterise typical musculoskeletal systems—kinetic energy, heat, or gravitational potential energy may serve as three illustrative examples. A convenient physical approach for such an analysis is to invoke the conservation of energy, which relates the work *W* that was done to the change Δ*E* in energy that results:

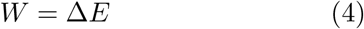

We use the conservation of energy to evaluate the flow of mechanical energy in the idealised model of a real musculoskeletal system that forms the basis of the theory of physiological similarity (Labonte 2023): a muscle with volume *V*_*m*_ exerts a force *F*_*i*_(*t*), and acts on a mass *m via* an anatomical arrangement with characteristic constant mechanical advantage *G*; there are no in-series elastic elements (Figure 1). The muscle can apply a force of no more than *F*_max_, shorten by a distance of at most *δ*_max_, and no faster than with a maximum shortening velocity 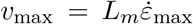, where *L*_*m*_ is a characteristic fascicle length, and 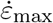 is a maximum strain rate in units muscle lengths per second. Throughout the initial analysis, it is assumed that the muscle can exert its maximum force until it has contracted by *δ*_max_, or until it has a achieved a shortening velocity *v*_max_, *F*_*i*_(*δ < δ*_max_, *v < v*_max_) = *F*_max_; above this maximum shortening distance and shortening speed, the force drops instantaneously to zero, *F*_*i*_(*δ > δ*_max_, *v > v*_max_) = 0. Implementing the force-length and force-velocity relationships as simple step functions is a simplification, and muscle does not behave in such an idealised way: the force it generates varies with both muscle length and muscle shortening speed before their limiting values are reached, *F*_*i*_(*δ >* 0, *v >* 0) *< F*_max_. We will demonstrate at the end of the results section that such more complex relationships considerably increase mathematical complexity without changing the general physical picture, so retrospectively justifying the idealisation.

To investigate how gearing can constrain the work output of muscle, we will first analyse the hypothetical case where muscle provides the only real force in the system. Muscle then only does work against the inertial force, the product between mass and the instantaneous acceleration. Each unit of muscle work *W*_*m*_ is thus entirely converted into kinetic energy, *K*:

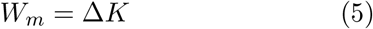

Although this scenario does not resemble any real system exactly, it is approximately correct as long as all external forces (*F*_*e*_) are much smaller than the output force of the musculoskeletal system, *F*_*e*_ *≪ F*_*o*_, which implies that *F*_net_ = *F*_*o*_ *− F*_*e*_ *≈ F*_*i*_*G*, and thus *W ≈ W*_*m*_. More importantly, it will focus the attention on the mechanism through which gearing can curtail muscle work output, and so help to understand the results of more complex analyses downstream.

To investigate how gearing influences the conversion of muscle work into different types of energy, we will next increase system complexity, and allow external forces *F*_*e*_(*t*) to oppose the motion of the mass. Such forces are “parasitic” in the sense that they do negative internal work, *W*_*e*_. The presence of external forces thus enforces a partitioning of each unit of muscle work into kinetic vs parasitic energy:

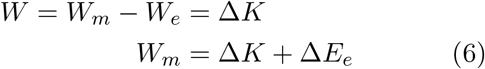

Where the analysis of an idealised inertial contraction was used to explain how gearing can curtail muscle work output, this step will be used to build a conceptual understanding of how gearing can influence the distribution of muscle work across kinetic and parasitic energy.

The results from both analyses will then be combined to determine an estimate of the mechanical advantage that provides the best compromise between the two opposing demands they revealed: for muscle to deliver its maximum work capacity, the mechanical advantage cannot be arbitrarily large; but to avoid this delivery taking diverging amounts of time, and to ensure that most of the muscle work flows into kinetic energy, *G* can also not be arbitrarily small.

In a last step, this result will be put to use to illustrate how “optimal” gearing depends on musculoskeletal anatomy, physiology and the mechanical environment in which the contraction takes place.

This demonstration will be conducted through the analysis of three simple case studies of real musculoskeletal systems.

## Results

### How can gearing influence muscle energy output?

For an inertial contraction starting from rest, conservation of energy implies:

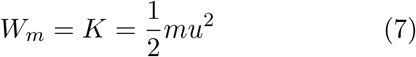

The problem is then reduced to the question of how much energy muscle can deliver. This question is as old as biomechanics itself (Borelli 1680), and the canonical answer is that the maximum energy output of muscle, its work capacity *W*_max_, is independent of the mechanical advantage. This assertion can be understood as follows: the work capacity of muscle depends on the maximum displacement-averaged force 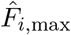 can exert as it shortens over a maximum contraction distance 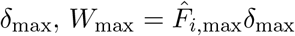. The work capacity of a geared muscle, in turn, is the product of the displacement-averaged external force 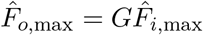 and the external displacement over which this force is moved, *x*_max_ = *δ*_max_*/G*. It follows that 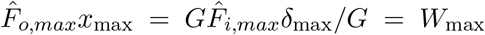 the work capacity of a geared muscle is the same as that of the ungeared muscle, and thus independent of the mechanical advantage.

If that was all there was to it, gearing would merely control how each unit of muscle work is split into the net average force, 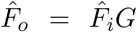 vs. the net displacement, *δ* = *xG*^*−*1^, but the muscle could deliver the same amount of work regardless of the value of *G*. Contractions would consequently always terminate at the same speed, the “Borelli-limit”, 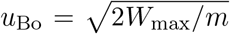 (Labonte 2023), and force-velocity trade-offs would only apply instantaneously but not ultimately (as suggested by McHenry 2010). But can the muscle deliver *W*_max_ for any value of *G*?

To identify the problem with this assertion, note that the output speed *u* is directly coupled to the muscle shortening speed *v, u* = *vG*^*−*1^. Inspect, then, the muscle shortening speed required to deliver the maximum work capacity for an arbitrary mechanical advantage, 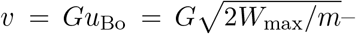 as *G* diverges, so must the muscle shortening speed. But muscle cannot shorten with arbitrarily large speeds. Instead, muscle shortening velocity has an axiomatic upper limit, *v* ≤ *v*_max_, so that the output speed is also bound by the “Hill-limit”, *u < u*_Hi_ = *v*_max_*G*^*−*1^ (Labonte 2023). This result is both elemental and crucial— muscle has not one, but *two* characteristic energy capacities. The work capacity, *W*_max_, is joined by a “kinetic energy capacity” (Labonte 2023; Labonte et al. 2024),

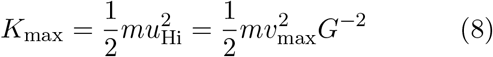

The work and the kinetic energy capacity pose two hard, independent bounds on muscle work output, *W*_*m*_ *≤ W*_max_ and *W*_*m*_ *≤ K*_max_, respectively. The maximum work muscle can deliver in a single contraction is consequently determined by whichever of the two limits is smaller (Labonte 2023) (Figure 2). Crucially, and unlike *W*_max_, *K*_max_ *∝ G*^*−*2^ depends explicitly on the mechanical advantage, and can thus be geared. As long as *K*_max_ *< W*_max_, gearing can therefore alter muscle energy output, and so affect the output speed that results from concentric muscle contractions (Labonte 2023; Labonte et al. 2024). In this regime, force-velocity trade-offs thus apply both instantaneously and ultimately, as argued by Arnold et al (Arnold et al. 2011).

It is convenient to determine an explicit relationship between mechanical advantage and muscle energy output in terms of the physical parameters of the musculoskeletal system. To find such a general condition, we first normalise eq. 7 by dividing both sides with *W*_max_, which yields a dimensionless form of the conservation of energy:

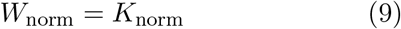

The crux is to recognise that both *W*_norm_ and *K*_norm_ are bound from above. The normalised energy output cannot exceed unity, *W*_norm_ *≤* 1 because muscle has a limiting work capacity; and the normalised kinetic energy cannot exceed 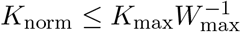, because muscle has a limiting kinetic energy capacity. The ratio of the two characteristic energy capacities, 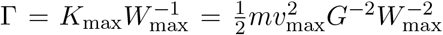, is the dimensionless “physiological similarity index”, so called because musculoskeletal systems that operate with equal Γ put out equal fractions of their maximal displacement, speed, work and power capacity in a single contraction (Labonte 2023).

Inspection of Γ permits direct identification of the dominant physical constraint on muscle energy output: if *G* is very large, then Γ *<* 1, and thus *K*_max_ *< W*_max_. The muscle is blocked from delivering its full work capacity, *W*_norm_ = Γ *<* 1, because it does not have a sufficiently high kinetic energy capacity— force-velocity trade-offs apply both instantaneously and ultimately (Figure 2). If *G* is very small, in turn, Γ *>* 1, and thus *K*_max_ *> W*_max_ —the muscle can deliver its full work capacity, *W*_norm_ = 1 *<* Γ, so that force-velocity trade-offs apply instantaneously, but not ultimately. The introduction of the physiological similarity index therefore reconciles the opposing views of McHenry and Arnold et al (Arnold et al. 2011; McHenry and Summers 2011)—both are correct, but in two opposite “design” regimes, characterised by small or large values of Γ, respectively. For an idealised muscle in an inertial contraction, the transition between both regimes occurs at Γ = 1, i. e. at the unique gear ratio for which the work and kinetic energy capacity of the musculoskeletal system are equal (Labonte 2023):

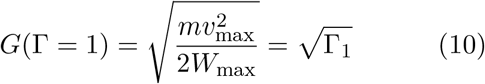

where Γ_1_ := Γ(*G* = 1) is the physiological similarity index for an “ungeared” muscle with a mechanical advantage of unity.

The key result of this first analysis may be summarised as follows: for muscle to deliver maximum energy, its kinetic energy capacity must be at least as large as its work capacity, Γ *≥* 1. Because the kinetic energy capacity can be geared, this condition can be achieved for any musculoskeletal system through appropriate gearing, 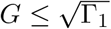 (Figure 2).

### How does gearing influence energy flow?

Consider next the more general case where muscle contracts against external forces, *F*_*e*_, which oppose motion. These external forces do negative work, and so redirect muscle work from kinetic into other forms of energy, *E*_*e*_ (Labonte 2023; McHenry 2012; Scholz et al. 2006). It is not immediately obvious how a fixed amount of muscle work will be split between *K* and *E*_*e*_. To determine this split, we re-write the energy balance (eq. 6)to find:

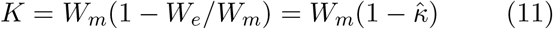

The key difference to the pure inertial case governed by eq. 7 is the dimensionless quantity 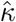, referred to as the “reduced parasitic energy” in the framework of physiological similarity (Labonte 2023). Without loss of generality, 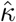 is the fraction of muscle work flowing into work against external forces. This fraction is determined by the ratio between the displacement-averaged opposing and driving force, respectively:

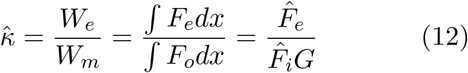

In the presence of external forces, it thus becomes important whether a unit of muscle work is delivered by displacing a large external force over a small displacement, or a small external force over a large displacement. The fraction of each unit of muscle work that flows into kinetic energy is maximised if it is delivered with diverging average force, 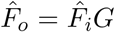, and vanishing displacement *x* = *δ/G*, i. e. for a diverging mechanical advantage.

To gain a more intuitive physical understanding of this mathematical result, consider a concrete example of two animals that seek to jump off the ground with a maximal takeoff velocity *u* (Figure 3). All else being equal, let one animal exhibit a smaller and the other a larger mechanical advantage. The muscles in both cases contract by the same amount, so that the work they deliver is identical, and equal to *W*_max_. However, the *external* movement that results from an equal amount of muscle shortening differs, and is larger for the animal with the smaller mechanical advantage. The work done against gravity is the product of this distance and the gravitational force, and is consequently larger, too. Because the total muscle work is identical across both cases, a difference in gravitational potential energy costs must imply a difference in the change in kinetic energy, Δ*K* = *W*_*m*_ *− W*_*g*_. Thus, the larger mechanical advantage results in both a larger average force *and* a larger take-off velocity, in defiance of the force-velocity trade-off; the height gained after take-off is higher for the jump with higher mechanical advantage.

**Figure 3:**
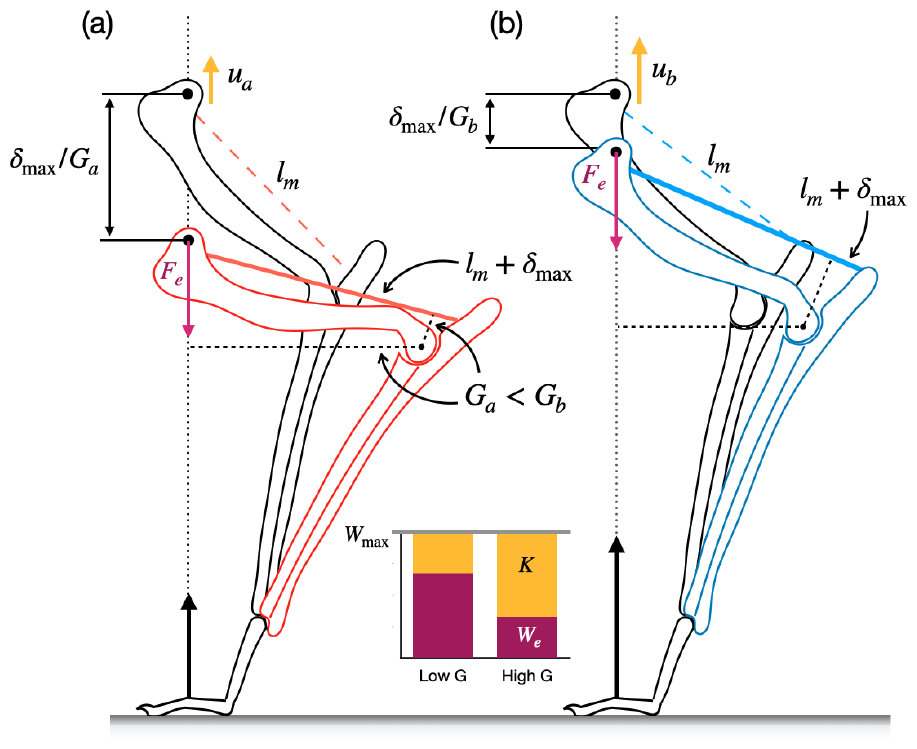
A thought-experiment with two jumping animals illustrates how the magnitude of the mechanical advantage, *G*, controls the partitioning of muscle work into kinetic energy, *K*, vs. parasitic work, *W*_*e*_; the morphology is based loosely on the mammalian forelimb. The two animals (a) and (b) are identical, except that (a) has a lower mechanical advantage at the elbow, *G*_*a*_ *< G*_*b*_. Let the muscles of both (a) and (b) deliver their maximum work capacity *W*_max_: the elbow extensors are first maximally stretched to a length *l*_*m*_ + *δ*_max_ (in colour), and then contract to accelerate the centre-of-mass until both animals have returned to their initial posture (in black). Because *G*_*a*_ *< G*_*b*_, animal (a) must crouch lower to stretch and contract its muscles by the same amount, so that the gravitational force must be moved over a larger *external* distance *δ*_max_*/G*_*a*_ during the acceleration phase. As a consequence, the work demanded to pay for the change in gravitational potential energy during the acceleration is larger in (a), *W*_*e*_ = *mgδ*_max_*/G*_*a*_, so that less work is available to increase the kinetic energy *K* = *W*_max_ *− W*_*e*_ (chart inset). Consequently, animal (b) achieves a higher takeoff speed *u*_*b*_ at the same takeoff height, and thus jumps higher above standing: a higher mechanical advantage results in a larger average net force, *and* a larger take-off velocity, in defiance of a force-velocity trade-off (see also Labonte 2023; Labonte et al. 2024; McHenry 2012; Scholz et al. 2006). This particular effect of *G* may be generalised for any external force through introduction of the reduced parasitic energy 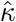, a dimensionless number which directly quantifies the fraction of muscle work lost to parasitic forces (see text for a mathematical definition).

The key result of this second step of the analysis may then be summarised as follows: movement is generally opposed by external forces, and these forces demand a share of muscle work; the work done by these external forces is equal to the product between their displacement-average, 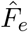, and the external displacement, *x*; and the mechanical advantage dictates the magnitude of the external displacement per unit muscle work. As a result, *G* controls the partitioning of muscle work into kinetic vs “parasitic” forms of energy.

### What constitutes optimal gearing?

It was demonstrated that gearing controls both the fraction of *W*_max_ that muscle can deliver, and how each unit of delivered work is partitioned into different forms of energy. For muscle to deliver *W*_max_, the mechanical advantage cannot be arbitrarily large; it must be small enough to ensure that *W*_max_ has been delivered exactly when or before *v*_max_ is reached. Although any mechanical advantage smaller than this critical condition will allow muscle to deliver *W*_max_, reducing *G* any further would bring two main disadvantages. First, it would reduce the average output force, so that delivering *W*_max_ will take longer—a musculoskeletal system operating exactly at the critical *G* delivers maximum work in minimal time, and thus maximises work and average power output simultaneously (Labonte 2023). Second, decreasing *G* also results in an increased loss of muscle work to parasitic forces, i. e. it reduces the transmission efficiency from work into kinetic energy, and consequently the output speed muscle can impart. There thus should exist an intermediate mechanical advantage that represents the optimal resolution of these competing demands (Labonte 2023; McHenry 2012; Olberding et al. 2019).

To find this “optimal” mechanical advantage, we first again obtain a dimensionless form of eq. 11. Division with *W*_max_ yields:

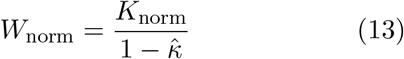

The optimal mechanical advantage of an idealised muscle is the critical value of *G* for which the maximum work capacity is delivered (*W*_norm_ = 1) exactly when the maximum shortening speed is reached (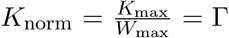 Figure 4). Eq. 13 can thus be written as:

**Figure 4:**
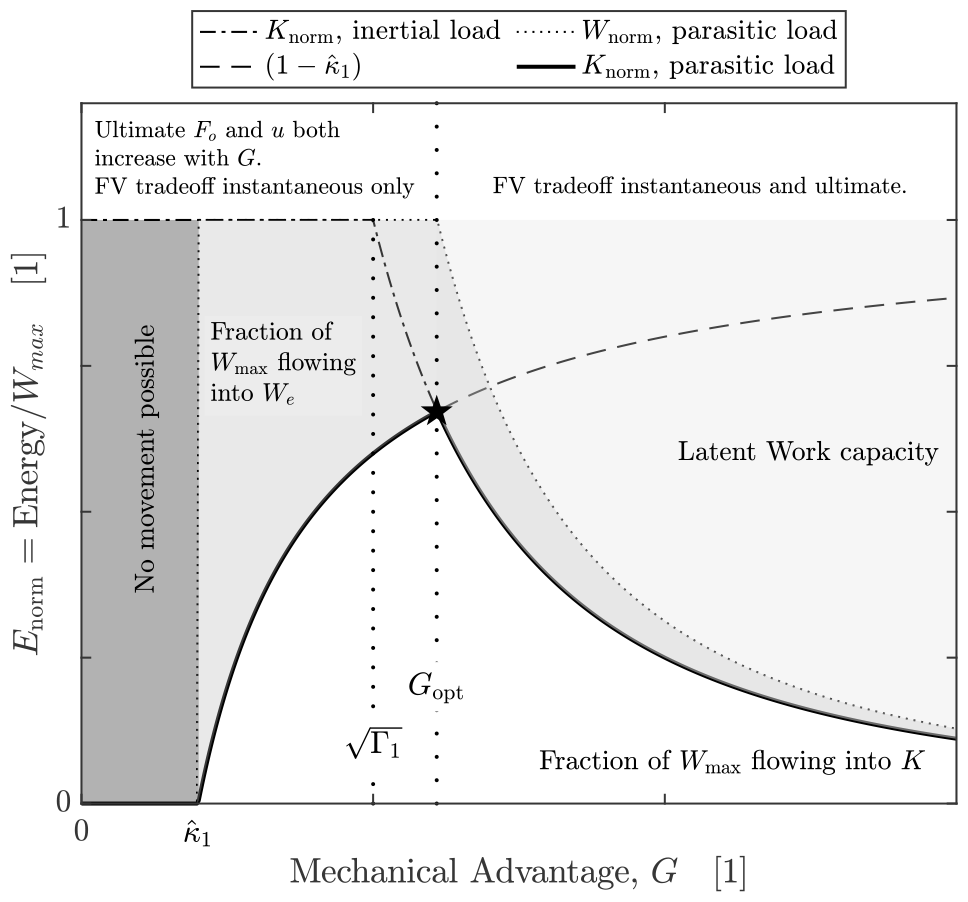
The energy landscape for an idealised musculoskeletal system in a contraction against a constant external force *F*_*e*_. The external force consumes a fraction 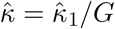 of the muscle work, and so alters the efficiency with which muscle work is transmitted into kinetic energy (Labonte 2023; McHenry 2012). If the mechanical advantage drops below a critical value 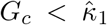, acceleration is no longer possible, as the muscle cannot resist the external force. For *G > G*_*c*_, *G* controls the fraction of the muscle work that flows into kinetic energy, 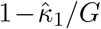. The optimal mechanical advantage *G*_opt_ (Eq. 15) now not only allows the system to deliver maximum work in minimal time, but also enables muscle to operate with maximal transmission efficiency; *G*_opt_ depends explicitly on Γ_1_ and 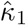, as defined by eq.15. For *G > G*_opt_, muscle work output is limited by its kinetic energy capacity. Note that here Γ_1_ = 1 and 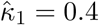 exact landscape appearance depends on the relative magnitude of Γ_1_ and 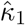, and is subtly different for parasitic forces that vary with speed, because equilibrium of forces now only occurs dynamically (see Figure 5f for an example)

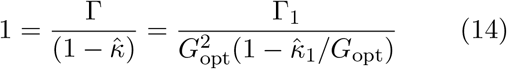

where we define 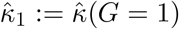 as the reduced parasitic energy of a musculoskeletal system with a gear ratio of unity. Eq. 14 is quadratic in *G*_opt_, and the positive root reads:

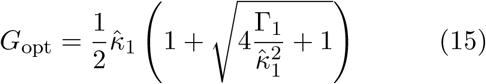

where we note that for all but the case of a constant external force, this expression is implicit, as 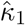 depends on *G* (see below for a more detailed discussion of this point).

The ratio 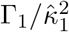 emerges as a key determinant of the optimal gear ratio. Two limiting cases may be considered. When 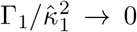, external forces are large relative to the system inertia; when 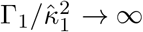, external forces are negligible. The optimal gear ratio in each limit follows as:

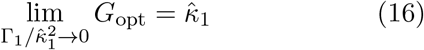

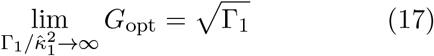

Both results may be interpreted as an equilibrium between the two dominant displacement-averaged forces (see also Labonte 2023; Richards and Clemente 2013b). When the external force is dominant, *G* is optimal when 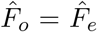 (for 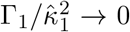)- and indeed, no motion is possible if ever 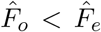. When inertial forces dominate, 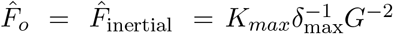 (for 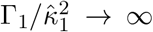). In many realistic scenarios, *G*_opt_ will fall between these two limits, and is defined by eq. 15.

The key result of this third step of the analysis may then be summarised as follows: *G* should neither be too large, for otherwise muscle can only deliver a fraction of its work capacity, nor can it be too small, for otherwise the time it takes to deliver each unit of work diverges, and most of it is lost to parasitic energy (Figure 4). These competing demands are encoded in two dimensionless numbers, which in combination define the optimal mechanical advantage: the value of *G* with which muscle can deliver maximum work in minimal time, and with maximum transmission efficiency.

Up to now, the analysis was agnostic to the nature of the external forces, and thus general; the aim was to build intuition for the key physical mechanisms at play, and to develop a theoretical framework that reveals the effect of gearing on the flow of mechanical energy within the system under study. We next apply this framework to three case studies to both illustrate its use and probe its explanatory power: the strike of a praying mantis, as an example for an approximately inertial contraction; the jump of kangaroo rat, as an example for a contraction against the constant gravitational force; and the leg stroke of a frog, as an example for a contraction against a velocity-dependent external force. We remind the reader of the simplifying assumption that the muscle force remains constant at its maximum throughout the contraction; this idealisation is discussed briefly and critically at the end.

### Case studies

Praying mantises exhibit a conspicuous adaptation of their forelimbs into a raptorial appendage, capable of catching prey in a predatory strike lasting less than 100 ms (Rossoni and Niven 2020). To catch prey effectively, they must be able to achieve maximum speed in minimal time—how should the raptorial appendage be geared to achieve this? Given the small appendage mass of around 0.17 g, and the vertical posture of the forelimbs (Figure 5a), the external forces are likely small compared to the muscle force, 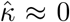. For an adult *Heirodula membranacea*, the strike occurs with Γ_1_ *≈* 4.4 *×* 10 ^*−*4^ (SI), and the optimal gear ratio follows as 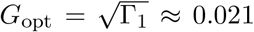, within the range of empirical estimates, *G*_emp_ *≈* [0.02, 0.05] (Figure 5b; Gray and Mill 1983).

**Figure 5:**
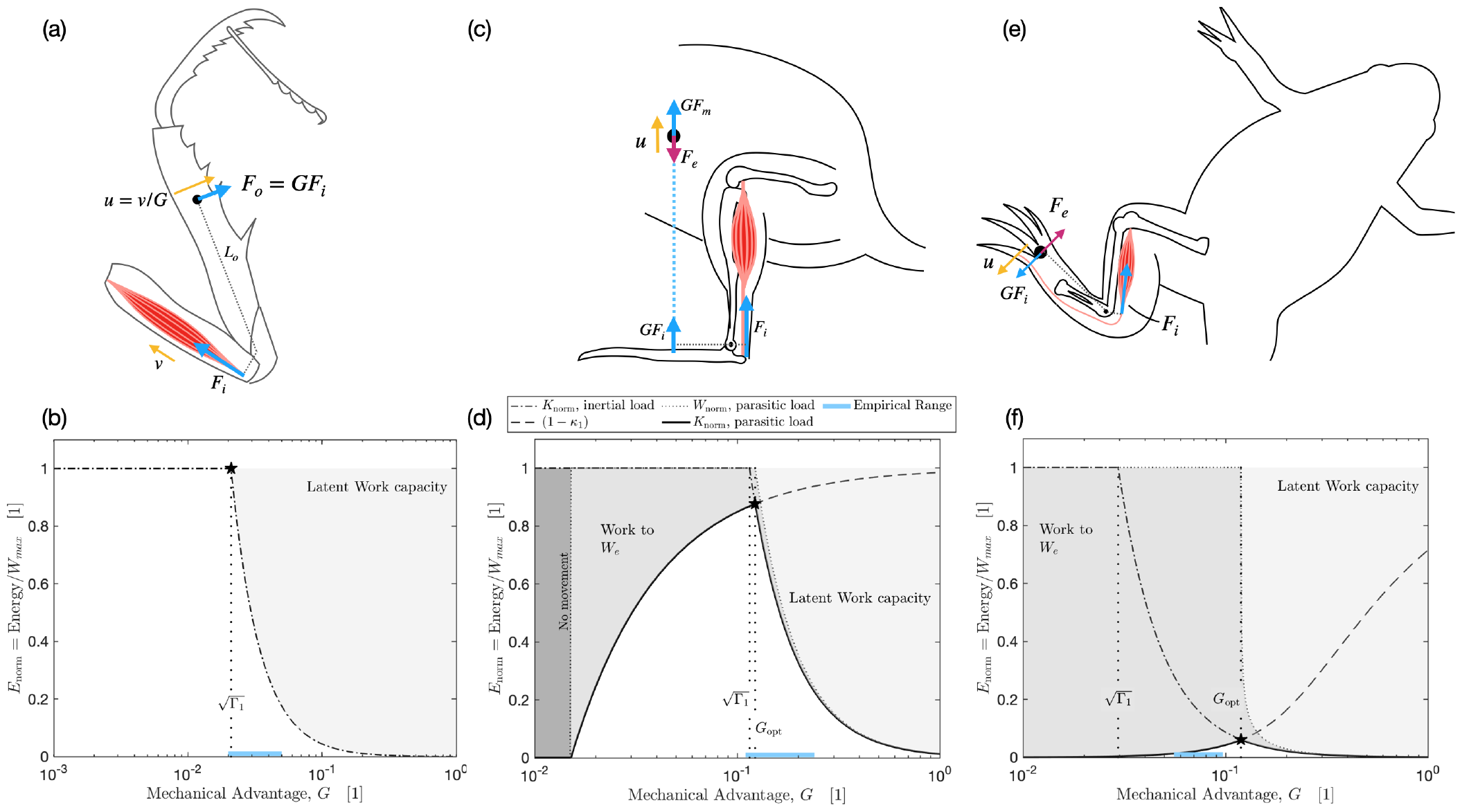
Three case studies exemplify how the mechanical environment influences the energy landscape for an idealised musculoskeletal system. In all cases, the optimal mechanical advantage, *G*_opt_, is defined as the value of *G* which enables the musculoskeletal system to deliver maximum work in minimal time, and with maximal transmission efficiency into kinetic energy (black stars). (a) The speed of a praying mantis strike is limited by the work muscle can do against an inertial load; external forces are negligible. It follows that 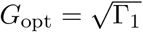 (black star). (b) A direct prediction falls within the range of empirical estimates for the mechanical advantage of the major coxo-femoral extensors in the mantis forelimb (blue line, see SI for details). (c) The takeoff jump speed of a kangaroo rat is determined by the work muscle can do against the inertial force and the gravitational force. (d) *G*_opt_ now depends on two dimensionless numbers, as defined by eq.15. A direct prediction (star) falls within the range of empirical values (blue line, see SI for details). (e) A frog swimming in a fluid medium contracts the plantaris muscle to push its foot rapidly through the water. (f) The empirical estimate for the mechanical advantage (blue line) is now far larger than the inertial optimum 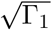, and close to the directly predicted optimum *G*_opt_ (black star, see SI for details). The energy landscape here differs from the gravitational case (panel d), as the external forces are velocity-dependent. As a result, there exists no minimal *G* below which movement is impossible. Furthermore, *W*_*e*_ is not “wasted” *per-se*, but a portion is transmitted to the body as kinetic energy. Due to the velocity-dependence of the external force, the mechanical advantage that maximizes foot kinetic energy also maximizes average and peak force.

The desert kangaroo rat is considerably larger (body mass of 106 g) and has hindlimbs adapted for quick escape jumps from predators, and serves as an example for a contraction against a non-negligible gravitational force, *F*_*e*_ = *mg*. Assuming that jumps are driven dominantly with the ankle extensors, we estimate Γ_1_ *≈* 0.013 (see SI)-about 30 times greater than for the mantis raptorial appendage. The external force is constant, so that the reduced parasitic energy follows as 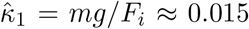. Eq. 15 provides an explicit condition for the optimal mechanical advantage, the value of *G* which maximises jump height above standing height, which yields *G*_opt_ *≈* 0.12; again within the range of empirical estimates, *G*_emp_ *≈* [0.11, 0.24]. Note that although the kangaroo rat is larger than the praying mantis, gravity still only plays a relatively minor role in determining the energy landscape, and the optimal mechanical advantage remains close to the inertial limit 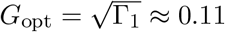.

Consider last the frog *Xenopus laevis*, which uses drag-based propulsion from its hind legs to swim through water. We seek to estimate the mechanical advantage of the plantaris that enables maximum foot speed. The contraction occurs with Γ_1_ *≈* 9 *×* 10^*−*4^ (see SI), and because the movement takes place in water, gravity can be neglected, but a drag force *F*_*D*_ must be considered. For *Xenopus* foot strokes, the drag force can be modelled as varying with the square of the speed, *F*_*D*_ = *β*_*Q*_*u*^2^ (*β*_*Q*_ is a constant quadratic drag multiplier of dimension kg m^*−*1^; Richards and Sawicki 2012)). Thus 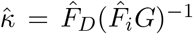, but as the displacement-averaged external force now depends on speed 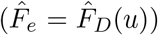, it also depends on *G*, so that eq. 15 is only implicit.

To the best of our judgement, no explicit and symbolic expression for *G*_opt_ exists for a quadratic drag force as a general function of external forces and Γ_1_ (see SI). However, the consideration of two limits is instructive. If the drag force is small compared to the inertial force, the optimal mechanical advantage approaches 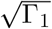 (Eq. 17); the system behaves approximately as if it was purely inertial. If the drag force is large compared to the inertial force, *G* is optimal if the muscle reaches its maximum shortening velocity exactly when the drag force and muscle output force *F*_*o*_ = *F*_max_*G* are equal (see SI). For the quadratic drag force,

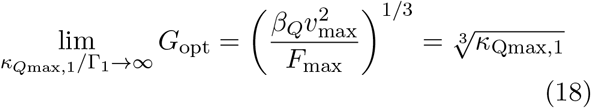

where *κ*_*Q*max,1_ is the ratio between the maximum ungeared drag and driving force, respectively. For the frog limb stroke, we estimate *κ*_*Q*max,1_ *≈*1.7*×* 10^*−*3^, so that *G*_opt_ = 0.03 in the inertial limit and *G*_opt_ *≈* 0.12 in the force limit. An explicit symbolic expression for the optimal gear ratio can be found for a linear drag force; this is shown in Figure 5f, where we note that the prediction for *G*_opt_ does not differ from Eq. 18 as the drag force is large. The empirical estimate of *G*_emp_ *∈* [0.06, 0.1] lies between these two limits, and the prediction from force balance is within 20% of the upper bound. We predict a maximal foot velocity of 1.2 m s^-1^, close to the empirical value of 1 m ^-1^ (Richards 2010).

No mathematical complexity should distract from the generality of the key result illustrated with these case studies: the mechanical advantage controls the fraction of the maximum work capacity muscle can deliver, because it determines the relative importance of the work and kinetic energy capacity; and it defines the fraction of each work which flows into kinetic energy, because it controls how work is partitioned between force and displacement. These two constraints are encoded by two unique dimensionless numbers: the ungeared physiological similarity index, Γ_1_, which depends explicitly on muscle physiology and musculoskeletal anatomy; and the ungeared reduced parasitic energy, 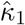, which reflects the influence of external forces and thus the physical characteristics of the specific environment in which the contractions takes place. The two dimensionless numbers can be combined to predict the magnitude of the optimal mechanical advantage for real musculoskeletal systems, and in three case studies, these predictions were either within or close to the range of empirical estimates.

### Further complexity: the force-velocity relationship of skeletal muscle

We have thus far analysed the flow of mechanical energy for a muscle with a force-velocity relationship idealised as a step-function: the muscle exerted maximum force for all shortening speeds up to its maximum shortening speed, *F*_*i*_(*v < v*_max_) = *F*_max_. A more realistic assumption is that the muscle force is maximal only for an isometric contraction *F*_*i*_(*v* = 0) = *F*_max_, and drops in some fashion with speed until it hits zero at the maximum contraction speed, *F*_*i*_(*v* = *v*_max_) = 0. How would this alter the key conclusions of this work?

The main investigative tool of this work has been the conservation of energy, 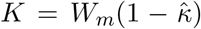. This conservation principle remains valid no matter the exact relationship between muscle force and velocity; it is completely general. As a consequence, the trade-offs that result in an optimal *G* must remain in place: if *G* is very large, muscle can only deliver a vanishing fraction of its maximum work capacity, and if *G* is very small, delivering any work takes a diverging amount of time, and the transmission efficiency goes to zero. However, the physical definition of *G*_opt_ is now more subtle, and determining its magnitude is significantly harder.

To understand the difficulty, note that the constraints *F*_*i*_(*v* = 0) = *F*_max_ and *F*_*i*_(*v* = *v*_max_) = 0 at once imply that the maximum work and kinetic energy capacity are now asymptotic constructs: they can be realised only in the limit of an infinitely slow (quasi-static) or infinitely fast (quasi-instantaneous) contraction, i. e. for Γ *→ ∞* and Γ *→* 0, respectively (Labonte 2023). As a result, the optimal mechanical advantage can no longer be defined as the value of *G* for which *K*_max_ and *W*_max_ are delivered simultaneously—no such contraction can exist. Instead, a conceptually equivalent *G*_opt_ may be defined as the mechanical advantage at which muscle delivers the most work in the least time (note well that this is distinct from requesting maximum average power, which is necessary, but not sufficient for *G* to be optimal Labonte 2023); to the best of our judgement, this requires numerical evaluation even for the most simple case of a linear force-velocity relationship (Labonte 2023).

Because *W*_max_ and *K*_max_ now only place an upper bound instead of providing an exact expression for the muscle energy output, the energy landscapes remain qualitatively, but not quantitatively identical (see SI for examples). For an inertial contraction, the difference between the idealised step-function and a more complex relation is maximal at Γ = 1, and depends in magnitude on the exact shape of the force-velocity relationship. For a linear force-velocity relationship, the achievable energy output deviates by no more than a maximum factor of two from the idealised step function for any value of Γ (see SI)—for many applications, this may be small enough to be considered negligible or within experimental error.

The influence of more complex force-velocity relationships must be restricted to small quantitative instead of broad qualitative differences, for the simple step function captures the key feature of the force-velocity relationship that introduces the fundamental physical principles at play: the muscle cannot generate a force for any *v > v*_max_. This is not to say that there are no interesting physical implications that emerge from a variation in force between 0 *< v < v*_max_. For example, the displacement-averaged muscle force now depends explicitly on *G*; the assessment of ultimate force-velocity trade-offs is thus complicated further. This and other subtleties surely warrant future investigation, but this exceeds the scope of this work. We will instead try to demonstrate how a focus on the broad physical effects of *G* on the flow of mechanical energy can provide a scaffolding for the systematical analysis of the comparative biomechanics of gearing in musculoskeletal systems.

## Discussion

Movement is integral to animal behaviour, can make the difference between life and death, and may demand the majority of an animal’s energy expenditure (Dickinson et al. 2000; Wilson et al. 2018, 2013). It therefore stands to reason that animal physiology and anatomy should bear the marks of selection for improved locomotor performance and efficiency. The mechanical advantage of musculoskeletal systems modulates the effect of muscle action on limb movement, and is thus one “design” element upon which selection may act. But what is the right mechanical advantage to select for?

An instantaneous analysis points to the force-velocity trade-off inherent in lever mechanics, and provides the basis for the most common functional interpretation of *G*—systems specialised for force should have large *G*, and systems specialised for speed should have small *G*. But recent work has demonstrated elegantly and unambiguously that instantaneous transmission does not necessarily map onto the ultimate outcome of a muscle contraction (McHenry 2010, 2012; Osgood et al. 2021). Inspired by this “energy perspective” on gearing, and aided by the theory of physiological similarity for muscle-driven motion (Labonte 2023), we have conducted a general mechanical analysis of the energy flow in a minimalist musculoskeletal system. This analysis yielded two key results. First, although gearing leaves the maximum work capacity of muscle unaffected, it can prevent muscle from delivering it, because it modulates the muscle’s kinetic energy capacity. Second, gearing also controls the partitioning of each unit of muscle work into kinetic vs parasitic energy, i. e. it influences the musculoskeletal transmission efficiency. For muscle to deliver *W*_max_, *G* must be sufficiently small; but for this work to be delivered rapidly, and to ensure that most of it flows into kinetic energy, *G* must be sufficiently large.

The relationship between the mechanical advantage and the achievable output speed is thus not a simple proportionality, as implied by the instantaneous perspective, but a complex function that depends on musculoskeletal anatomy, physiology, and the physical environment in which the contraction takes place (McHenry 2010, 2012).

Where the instantaneous perspective enables a functional interpretation of the mechanical advantage only through direct comparison—system *A* has a lower *G* than system *B*, and has therefore been selected for speed—the theory of physiological similarity permits a *direct* estimation of the specific mechanical advantage that allows musculoskeletal systems to deliver *W*_max_ in minimal time, and with maximal transmission efficiency. In musculoskeletal systems that have been selected for dynamic movements, this mechanical advantage may reasonably be considered “optimal” (Labonte 2023; Olberding et al. 2019). In the following discussion, we will put this optimality criterion to work to analyse three different aspects of the functional anatomy of musculoskeletal systems.

First, we will investigate what the theory of physiological similarity reveals about the magnitude of the optimal mechanical advantage, and show that it provides a speculative explanation for why the mechanical advantage of the vast majority of musculoskeletal systems is smaller than unity. Second, we will assess the variation of the optimal *G* with environment and animal size, and present an mechanical energy-centred hypothesis for why terrestrial animals of different size tend to vary in their posture. And third, we will discuss whether an energy perspective renders the use of *G* as simple proxy for selection for speed vs force obsolete, as argued forcefully by McHenry (McHenry and Summers 2011; McHenry 2012). It will be demonstrated that, even if many musculoskeletal systems occupied a mechanical regime where a reduction in *G* increases muscle work output, as argued by Arnold et al. (2011) in their critical response to McHenry, there exists an architectural “many-to-one-mapping”, such that covariation of independent geometric variables can ensure physiological similarity. That is, the mechanical performance of two musculoskeletal systems with equal muscle volume, payload and physiology can be identical even when *G* differs.

### Borelli’s riddle, and the magnitude of the optimal mechanical advantage

*Who indeed would be stupid enough to [*…*] use a machine or contrivance not to save forces but rather to spend forces?*

Functional interpretations of *G* are typically grounded in an instantaneous perspective: specialisation for speed requires small *G*, and specialisation for force requires large *G*. Irrespective of whether this interpretation is sound, it has two shortcomings: it reveals nothing about the actual magnitude of a functionally optimised mechanical advantage; and, consequently, it can only ever allow functional interpretation through comparison. It was first observed by Borelli (Borelli 1680), and has been confirmed many times since, that the mechanical advantage of most musculoskeletal systems is smaller than unity, i. e. gearing usually amplifies displacement and velocity, and thus attenuates force. But why should evolution be “stupid enough” to select systems with small mechanical advantage (Borelli 1680; Pope 2005)?

To answer “Borelli’s riddle”, we posit that selection should often favour mechanical advantages that enable muscle to deliver its maximum work capacity in minimal time, and that ensure that most of this energy is converted into kinetic energy. In such a system, *G* can neither be too large, for muscle would then only have access to a fraction of *W*_max_, nor can it be too small, for otherwise it would take longer to deliver *W*_max_, and an increasing amount of muscle work would flow into parasitic instead of kinetic energy (Labonte 2023). The “optimal” mechanical advantage should thus be of intermediate magnitude. We have shown that this magnitude can be expressed in terms of two characteristic dimensionless numbers: the physiological similarity index, Γ_1_, and the reduced parasitic energy, 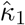 (eq. 15). Because these numbers are formed from the physiological and anatomical properties of the musculoskeletal system, the pay load, and the physical environment in which the contraction takes place, there exist complex interdependencies, and the magnitude of *G* alone says little about mechanical performance.

To investigate what a focus on mechanical energy reveals about Borelli’s riddle, we will estimate the optimal gear ratio for the simplest case: a muscle contraction against an inertial load, i. e. all external forces are of negligible magnitude. Although this may sound artificial, it is likely applicable to a large range of musculoskeletal systems, for the muscle force is much larger than the weight force in all but the heaviest vertebrate animals (Alexander 1985; Labonte 2023; Labonte et al. 2024).

In an inertial contraction, a muscle will deliver maximum work in minimal time if the ratio between its kinetic energy and work capacity is unity (Labonte 2023). For the minimalist musculoskeletal system studied in this work, this condition can be written as:

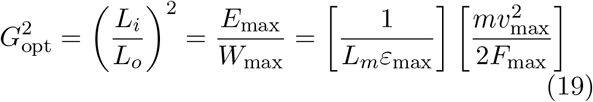

where *ε*_max_ is the maximum strain. The right-hand side was written as the product of two fractions to illustrate that the mechanical advantage is optimal if the squared ratio of the two anatomical lengths that define it is equal to the ratio between two physiological lengths: the maximum muscle shortening distance, *L*_*m*_*ε*, and the distance over which an ungeared muscle would have to shorten to accelerate to its maximum shortening speed, *mv*^2^(2*F*)^*−*1^. The mechanical advantage effectively acts as a modulator of this second term, such that an appropriate choice of *G* ensures that the two length are equal at the system level. In order to provide a rough estimate of *G*_*opt*_, we find that for a “typical” striated muscle (see SI):

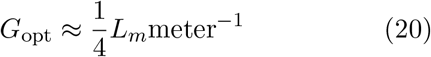

Thus, the characteristic muscle fascicle length *L*_*m*_ would have to exceed 4 m to demand *G*_*opt*_ *>* 1, providing a speculative answer to Borelli’s riddle, grounded in first principles: the mechanical advantage of most musculoskeletal systems “spends forces”, i. e. is smaller than unity, because larger gear ratios would constrain muscle work output to be sub-maximal.

Because *G* is a geometric parameter than can help to navigate physiological constraints, we suggest to interpret it as an anatomical adaptation that relaxes selective pressure on muscle anatomy: through appropriate gearing, optimal performance can be achieved for any value of *L*_*m*_. For a fascicle length of 10 cm, *G*_opt_ *≈* 0.03, and for *L*_*m*_ = 1 cm, it drops to *G*_opt_ *≈* 0.003—although the fascicle length differs by a factor of ten, both systems can achieve equivalent mechanical performance, provided that the magnitude of Γ is identical, so that they are physiologically similar (Labonte 2023); gearing provides an architectural degree of freedom in musculoskeletal “design”. The strong size-dependence of *G*_opt_ is apparent, and the focus of the next section.

### Osborn’s conjecture, and the size-dependence of the optimal mechanical advantage

*The straightening of the limb [in large quadrupeds] is an adaptation designed to transmit the increasing weight through a vertical shaft*.

Henry Fairfield Osborn (Osborn 1900)

The optimal mechanical advantage for an idealised muscle is determined by the interplay of two dimensionless numbers: the ungeared physiological similarity index, Γ_1_, defined by the physiological and anatomical properties of the muscle and the payload; and the ungeared reduced parasitic energy, 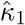, which depends also on the specific physical environment in which the contraction takes place. How do both numbers vary with animal size?

Under the parsimonious assumptions of isogeometry and isophysiology, Γ_1_ *∝ m*^2*/*3^ (Labonte 2023). The scaling of 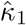 in turn depends on the nature of the external force. For the gravitational force, of relevance for terrestrial locomotion, isogeometry implies 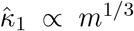 for a linear drag force one may find 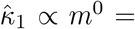 constant; and for a quadratic drag force, 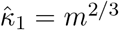. It follows at once that *G*_opt_ should vary systemically with size, and that the details of this variation should depend on the physical environment. Thus, and as is increasingly recognised for other musculoskeletal “design” elements such as the stiffness of in-series elastic elements (Galantis and Woledge 2003; Ilton et al. 2018; Lichtwark and Wilson 2008, 2005; Mendoza and Azizi 2021; Rosario et al. 2016), optimal gearing requires “tuning” to other physical and anatomical properties (Olberding et al. 2019). Is there any evidence that selection has “tuned” *G* to size and environment, as required for an optimisation of mechanical energy flow?

It has long been noted that larger mammals tend to adopt a more upright posture than smaller mammals (Gray 1944; Gregory 1912b; Osborn 1900). It was pointed out by Biewener (Biewener 1989b) that this postural variation brings about a systematic variation in the “effective mechanical advantage” (EMA) of the limbs: crouching increases the moment arm of the ground reaction force with respect to the limb joints, and thus reduces the mechanical advantage averaged across all limb muscles. We extracted empirical data for EMA for 16 mammalian quadrupeds from the literature, and estimate Γ_1_ *≈* 0.0063*m*^2*/*3^kg^*−*2*/*3^ and 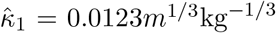 (see SI). Eq. 15 then yields *G*_opt_ *≈* 1*/*11mass^1*/*3^, compared to EMA = 1*/*4*m*^1*/*6^, estimated via an ordinary least squares regression on log10-transformed data (Fig. 6. We note that a regression restricted to Biewener’s original data, yields EMA *≈* 1*/*4*m*^0.24^).

**Figure 6:**
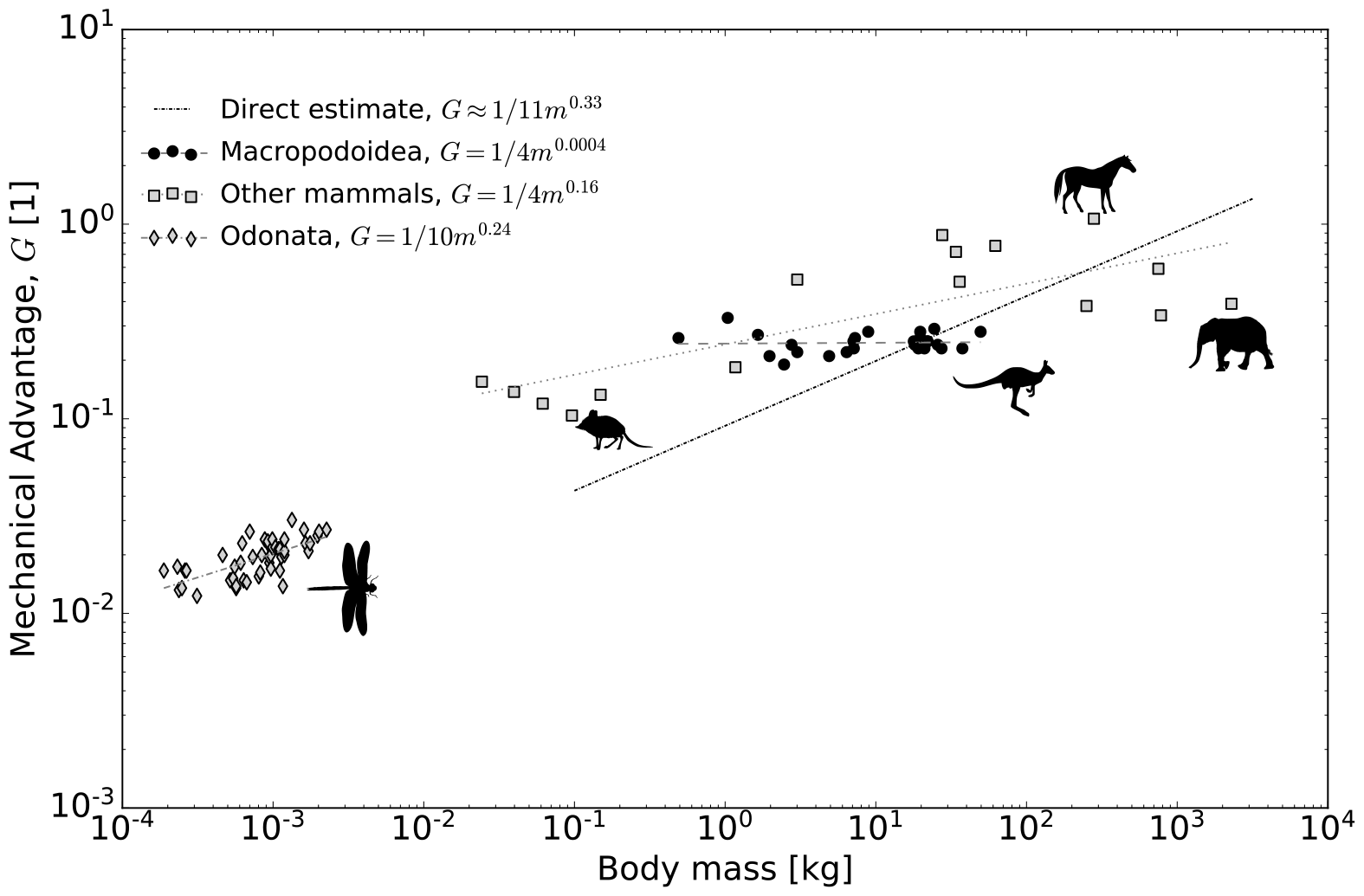
The optimal mechanical advantage is determined by two dimensionless numbers: the ungeared physiological similarity index, Γ_1_, which encodes the competition between kinetic energy and work capacity, and the reduced parasitic energy, 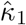, which encodes the relevance of external forces compared to the driving force. Because both numbers vary with body mass *m* under the parsimonious assumptions of isogeometry and isophysiology, so does the optimal mechanical advantage. For a contraction against the gravitational force, the inertial and the parasitic limit yield the same scaling coefficient, 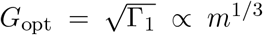, vs 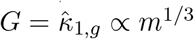, respectively. As an illustrative example, we directly predict the magnitude of this optimal mechanical advantage for quadrupeds (dark dotted line), and compare it to the result of an ordinary least squares (OLS) regression on the effective mechanical advantage for 16 species covering about six orders of magnitude in body mass (see (Basu and Hutchinson 2022; Biewener 1989b, 2005; Ren et al. 2010) and SI for details). For a contraction against a quadratic drag force, the scaling coefficient for *G*_opt_ should fall between the parasitic limit, 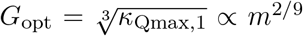, and the inertial limit 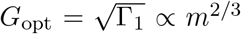. As an illustrative example, we extracted the mechanical advantage of the basalar flight muscle for eight dragonfly species (see Schilder and Marden (2004) and SI for details). OLS regression yields a slope of 0.24, in close agreement with the theoretical estimate. Note that *G* may also be independent of size, as occurs in Macropodoidea (Bennett and Taylor 1995).

In notable resemblance of the scaling of EMA in terrestrial mammals, the mechanical advantage of the basalar muscle in eight species of dragonflies increases with mass as *G* = 1*/*10*m*^0.24^ (Schilder and Marden 2004). We were unable to extract estimates for all physical quantities required to directly predict *G*_opt_, and thus only evaluate the scaling.

Dragonflies fly at Reynolds numbers Re *>* 10^3^, so that the drag force likely varies with the square of the speed. *G* should thus scale somewhere between *G ∝ m*^2*/*9^ = *m*^0.222^ and *G ∝ m*^1*/*3^, in robust agreement with the empirical data, *G ∝ m*^0.24^ (SI, Fig. 6).

This initial evidence for “tuning” is partially encouraging, and thus partially concerning. It is encouraging, for theoretical predictions are not only in qualitative, but also in a fair degree of quantitative agreement with empirical data: the elevation for the EMA for terrestrial mammals is within a factor of three of the empirical data, and the scaling of the mechanical advantage of dragonfly flight muscle is almost identical to the prediction, and certainly within the confidence intervals (see SI). In light of the simplicity of the analysis, and the many real features of musculoskeletal systems it ignored—muscle pennation, variations in *G* throughout the contraction, in-series elastic elements, and a plausible dependence of the maximum muscle strain on *G*—the extent of this agreement at the very least fails to reject the hypothesis that the identified mechanical constraints may hold some sway over musculoskeletal design.

However, it is also clear that a simple mechanical prediction does not tell the whole story, for the slope of EMA in terrestrial mammals deviates meaningfully from it; indeed some animal groups such as macropods show no size-specific variation in EMA at all (Fig. 6 and (Bennett and Taylor 1995; Thornton et al. 2024)). Larger terrestrial mammals are also generally faster, and eventually slow down; both findings suggesting that gearing may not be “optimised” to keep energy output and transmission efficiency independent of animal size (Labonte et al. 2024). Perhaps this deviation arises because some key mechanical constraints were omitted in the analysis. For example, a scaling of EMA *∝ m*^1*/*3^ may keep the muscle forces required to ensure static equilibrium at bay (Biewener 1989b), but it will result in a positive allometry of the peak ground reaction force during dynamic contractions, and so likely reduce bone safety factors. And perhaps some discrepancy should not be surprising altogether, for animal locomotion is constrained by more than just mechanics, and a holistic integrative perspective also ought to consider locomotor economy and neuromechanical control. For now, the influence of *G* on the flow of mechanical energy provides at least one further hypothesis not only for why larger animals should operate with larger *G* (Biewener 1989b), but also for why smaller animals should operate with smaller *G* (Usherwood 2013).

### McHenry’s objection, and the functional significance of the mechanical advantage

*This is a message [*…*] that challenges an assumption implicit in a century of literature*.

Matthew McHenry and Adam Summers (McHenry and Summers 2011)

By demonstrating that a variation in *G* is neither necessary nor sufficient to alter the maximum speed a musculoskeletal system can impart, McHenry cast doubt on the widespread practice to use *G* as a proxy for functional specialisation for force vs velocity (McHenry and Summers 2011; McHenry 2012)— an objection that has been echoed by others since (Osgood et al. 2021). In the forward simulations that supported the argument, the energy input was fixed, so that, without internal or external dissipative forces, force-velocity trade-offs could only ever apply instantaneously. But can muscle deliver the same energy regardless the value of *G*?

By analysing the flow of mechanical energy during contractions with arbitrary *G*, it was demonstrated that gearing alters the musculoskeletal kinetic energy capacity, and in doing so can curtail muscle work output even in the absence of dissipative forces. It follows that that there is large design space where a smaller value of *G* may well be associated with a lower force and a larger maximum output velocity— the classic force-velocity trade-off. Although there are few if any musculoskeletal system where robust information on all relevant parameters is available, rough estimates suggest that many systems may fall into this parameter space, i. e. 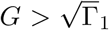 (see above and Labonte 2023; Labonte et al. 2024). Does this mean that the instantaneous perspective is salvaged, and that *G* can continue its service as a simple anatomical index for musculoskeletal specialisation for force vs speed?

It was already noted above that the optimal mechanical advantage is size- and environment-dependent—this result in and of itself sends a clear warning against a functional classification of musculoskeletal systems on the basis of *G* alone. However, matters get worse still, for it will now be shown that muscle mechanical performance also depends on other geometric properties of the musculoskeletal system. In other words, even for two musculoskeletal systems that operate in the same environment, are of the same size, have the same muscle volume, and identical muscle physiology, the magnitude of *G* does not provide a reliable indication for specialisation for force vs velocity.

To put this assertion to the proof, we relate the fascicle length to the muscle volume and cross-sectional area, *L*_*m*_ = *V*_*m*_*/A*_*m*_, introduce the muscle aspect ratio, 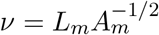, and re-write eq. 19 to find:

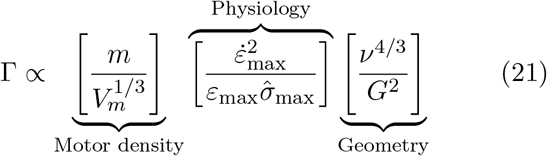

where the right-hand side was written as the product of separate terms to highlight three distinct components of the physiological similarity index (Labonte 2023): a ratio between pay load mass and motor size, muscle physiology, and musculoskeletal geometry. The motor density encodes a size-dependence for isogeometric systems, Γ *∝ m*^2*/*3^; the physiology is parsimoniously considered conserved; but the geometry term reveals a major problem for the typical comparative interpretation of *G*: systems with different *G* may well be physiologically similar, that is have equal Γ (Labonte 2023), even if neither muscle physiology nor investment differ—variations in the muscle aspect ratio *ν* can cancel the effects of variations in *G* (Figure 7). It is unclear why the ratio between in- and out-lever should possess a higher evolutionary flexibility than the muscle aspect ratio—both are geometric parameters defined by the ratio of two characteristic lengths. Thus, *even for a system in which a variation in G may modulate muscle energy output, G cannot be reliably interpreted as a sign for selection for force vs speed* —neither instantaneously, nor in terms of mechanical energy output. Instead, robust functional interpretation requires to also consider at least the muscle aspect ratio, but *ν* is rarely reported or discussed in comparative studies on the mechanical advantage.

**Figure 7:**
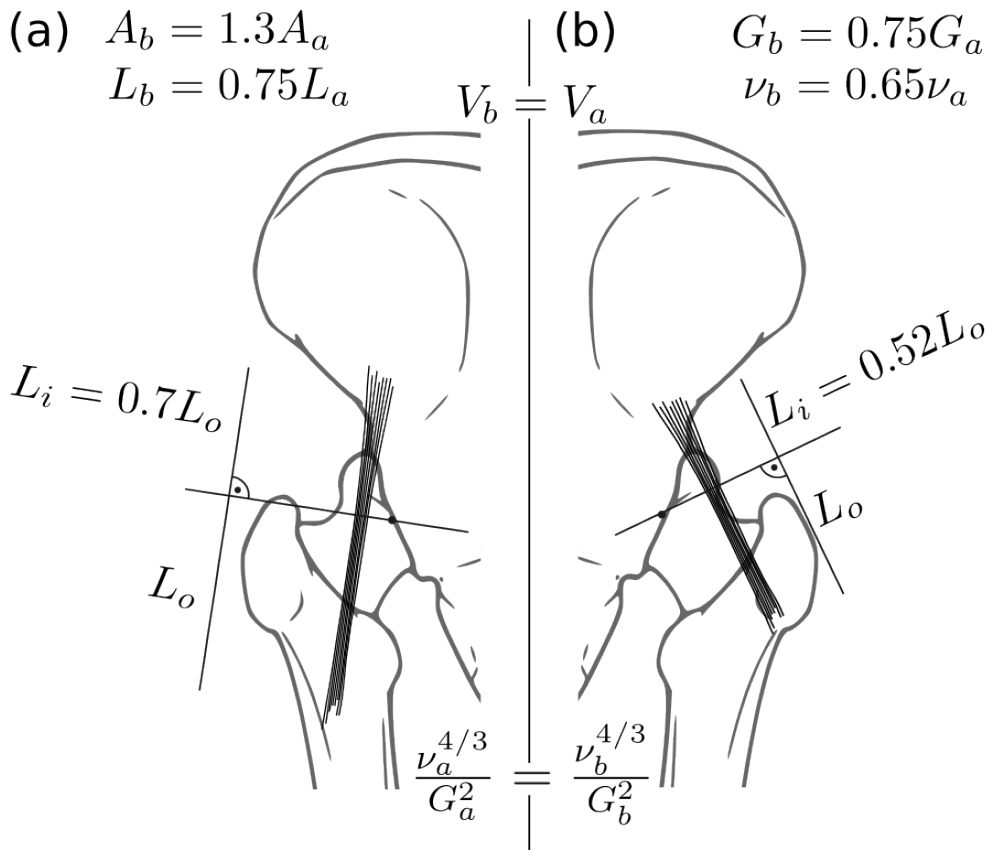
Variations of the muscle aspect ratio, 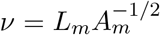 can “cancel” the effect of variations in the mechanical advantage *G*, in the sense that musculoskeletal systems with different *G* may nevertheless operate with equal physiological similarity index Γ *∝ ν*^4*/*3^*G*^*−*2^. As an illustrative example, consider two different geometrical arrangements of the Gluteus medius hip abductor; muscle volume and physiology are identical, but the mechanical advantage differs by a factor *G*_*a*_*/G*_*b*_ = 1.45. As long as Γ *<* 1, and all else being equal, a change in *G* will result in instantaneous and ultimate force-velocity trade-offs. But it would nevertheless be premature to conclude that (a) is specialised for force, and (b) is specialised for speed: the muscle in (b) has a larger cross-sectional area, and shorter fascicle length, so that both arrangements operate with equal Γ. Because of this many-to-one-mapping, differences in mechanical advantage between study organisms cannot serve as reliable indicator for specialisation for force vs velocity by itself—neither instantaneously, nor ultimately.

On the basis of these arguments, we join McHenry and others in their concern that an instantaneous interpretation of *G* can be functionally misleading (McHenry and Summers 2011; McHenry 2012); great care must be taken in resting the functional analysis of skeletal anatomy on a comparison of the mechanical advantage alone. Even in systems that are “overgeared”, so that instantaneous force-velocity trade-offs can map onto contraction outcomes as suggested by Arnold et al (Arnold et al. 2011), independent variation in geometric variables such as the muscle aspect ratio can result in the mapping of many geometric designs onto the same mechanical performance: musculoskeletal systems with markedly distinct geometries can be physiologically similar (Labonte 2023). Such “many-to-one-mapping” has been identified in an increasing number of mechanical systems (e. g. Alfaro et al. 2005; Arnold 1983; Blanke et al. 2018; Chatar et al. 2022; Moen 2019; Strobbe et al. 2009), and can enable morphological and physiological diversification despite a convergence in mechanical function (Muñoz 2019; Muñoz et al. 2018; Wainwright 2007; Wainwright et al. 2005). Viewed in this light, our warning against an isolated interpretation of *G* is not a pessimistic negation of the functional significance of *G*, but an optimistic proposal to re-evaluate the anatomical diversity of musculoskeletal anatomy through the functional interpretation derived from the theory of physiological similarity.

### Outlook

A mechanical analysis of a minimalist musculoskeletal system suggested a physical interpretation of the mechanical advantage that is robustly rooted in the conservation of mechanical energy. Illustrative applications of this framework suggested promising qualitative and semi-quantitative agreement between theoretical predictions and observable features of musculoskeletal systems. However, this initial success must not be allowed to mask the fact that several essential aspects that add further complexity were omitted: the mechanical advantage does not remain constant throughout a contraction Carrier et al. (1998b, 1994); Olberding et al. (2019); elastic elements such as tendons, apodemes and aponeuroses may obscure the relationship between muscle input and skeletal output (Eng and Roberts 2018; Galantis and Woledge 2003; Olberding et al. 2019); we only provided a cursory treatment of the effect of force-velocity properties, and neglected force-length properties; many muscles are pennate, and the pennation angle thus provides another architectural degree of freedom; gearing due to pennation can depend on the magnitude of the applied muscle force (Azizi et al. 2008; Eng et al. 2018); and this relation between force magnitude and gearing likely varies with other key biological variables such as age (Holt et al. 2016). Without doubt, more complex and accurate mechanical analyses are in order to put the merits of the theory of physiological similarity to a rigorous test; the problem requires further attention, and many details remain unresolved.

The combination of mathematical simplicity, experimental accessibility, and functional consequence has rendered *G* a rather attractive anatomical metric. These are no doubt desirable qualities of scientific theories, but perhaps an exclusively instantaneous interpretation is so simple that it risks to blind us to other relevant functional implications of variations in *G*. Analysing the effects of gearing in terms of mechanical energy permits a more holistic analysis of the functional and comparative anatomy of musculoskeletal systems than an instantaneous perspective affords, and brings about the exciting opportunity to integrate mechanical analyses with powerful evolutionary concepts such as many-to-one-mapping, morphological modularity and integration. Such future work holds significant promise to progress our understanding of the evolutionary biomechanics of animal movement.

## Acknowledgments

This study was supported by the Curiosity Fund, generously awarded to DP and DL by themselves. DL thanks Chris Clemente, Natalie Holt, Chris Richards and Jim Usherwood for many insightful discussions about gearing. DP thanks the OSyM network for the invitation and support to present this research at SICB and contribute this paper.

## Supplementary Information

### Optimal gearing for velocity-dependent external forces

#### General procedure

The external force *F*_*e*_ may depend on the velocity, *F*_*e*_(*u*). A common biological example of velocity-dependent external force is the drag force, *F*_*D*_. Drag is a complex physical phenomenon, and a first-order model typically considers a drag force that varies either linearly or quadratically with speed. A linear drag force, *F*_*D*_ *∝ u*, is an appropriate model for viscous drag, which dominates at small Reynolds numbers (that is, slow speed, high viscosity, or small length). At intermediate to large Reynolds numbers, a quadratic drag force, *F*_*D*_ *∝ u*^2^, is more appropriate.

The general analysis for both a linear and quadratic drag force proceeds from the conservation of energy:

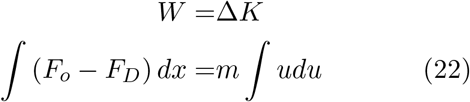

Because the drag force depends on *u*, solution requires separation of variables:

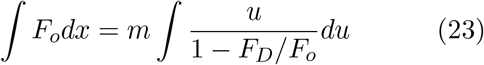

To introduce the mechanical advantage, we use the couplings *F*_0_ = *F*_*i*_*G*, and *x* = *δ/G*, and note that *G* is optimal if the maximum displacement and shortening speed of muscle are reached simultaneously:

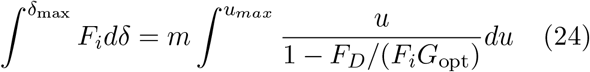

For a linear drag force, *F*_*D,L*_ = *β*_*L*_*u*, the velocity integral evaluates to:

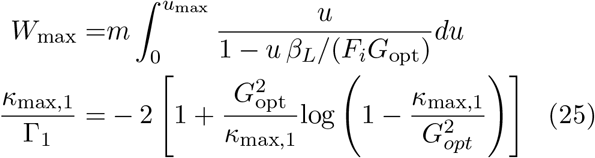

where we used the coupling *u* = *v/G*, and introduced 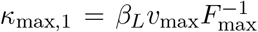 as the ratio of the maximum drag and driving force for an ungeared muscle with a mechanical advantage of unity.

For a quadratic drag force, *F*_*D,Q*_ = *β*_*Q*_*u*^2^, one finds instead:

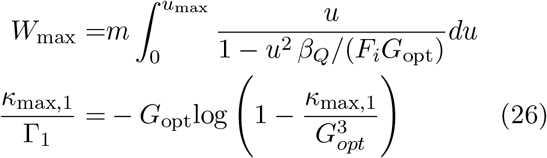

where 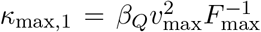 has the same physical meaning as before.

Thus, for both a linear and a quadratic drag force, the optimal mechanical advantage is determined uniquely by two dimensionless numbers, Γ_1_ and *κ*_max,1_.

#### The optimal mechanical advantage for linear drag

For the linear drag force, an explicit solution for *G*_*opt*_ can be found:

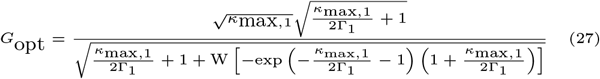

where *W* is the Lambert W function. It is hard to develop an intuitive feel for this expression, but the limits provide a clear physical picture. For convenience, we define *γ* = *κ*_max,1_*/*(2Γ_1_), and consider the limits *γ →* 0 (inertial forces dominate) and *γ → ∞* (external force dominate). Through Taylor expansion about *γ* = 0, we find lim_*γ→*0_ 1 + W [*−*exp (*−γ −* 1) (1 + *γ*)] = *γ*. The right-side limit for the remaining term is:

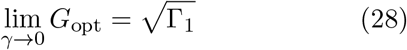

In this limit, inertial forces dominate, and optimal mechanical advantage is thus independent of external forces. For *γ → ∞*, the productlog-term goes to 0, which yields:

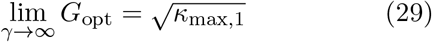

The inertial force is now irrelevant, and the optimal mechanical advantage is the value of *G* which ensures an equilibrium of the parasitic and driving force.

For any intermediate value, the optimal gear ratio depends on a complex combination of the three key forces: the inertial force, the driving force, and the parasitic force.

#### The optimal mechanical advantage for quadratic drag

To the best of our judgement, the implicit expression for *G*_opt_ for a quadratic drag force allows no explicit writing. Instead, *G*_opt_ has to be determined numerically. However, the limits can still be assessed, and follow in direct analogy to the case for a linear drag force.

When the drag force is large, *γ* = *κ*_max,1_*/*(Γ_1_) *→ ∞*, and eq. 26 yields:

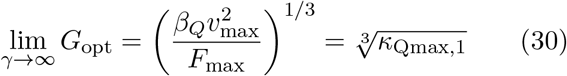

which is equivalent to equilibrium of maximum dynamic forces (see also Richards and Clemente (2013b)):

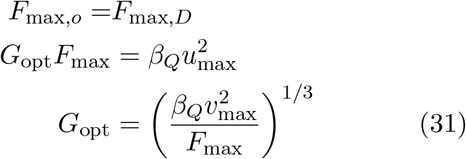

When inertial forces dominate, 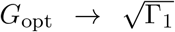, which follows as before.

Recognizing that both quadratic and linear drag have the same inertial limit for *G*_opt_, the symbolic results for linear drag above can be used to approximate quadratic drag by setting the upper *G*_opt_ limits to be equal. This implies:

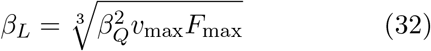

### Energy flow of geared muscle with force-velocity properties

Real muscle has more complex force-velocity properties than the idealised muscle studied in most of this work: the force it can exert is sub-maximal for any shortening speed larger than zero. For a “Hill-muscle”, the force varies between a maximum *F*_*i*_(*v* = 0) = *F*_max_ in an isometric contraction, and a minimum *F*_*i*_(*v* = *v*_max_) = 0 during a contraction with maximum speed. By approximating the relationship between these two extremes as linear, explicit symbolic expressions for the energy landscape can still be derived, provided that the external force is constant (Labonte et al. 2024). The quantitative appearance of the energy landscapes clearly differs between an idealised and a Hill-muscle (Fig.8), but the qualitative picture remains unchanged: muscle work output vanishes as *G* diverges; and as *G* vanishes, delivering the same amount of work takes more time, and the fraction that flows into kinetic energy, the transmission efficiency 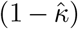, approaches zero– the same trade-offs exist, and the optimal mechanical advantage will be of intermediate magnitude. This result arises, because the idealised and the Hill-muscle become indistinguishable at the limits corresponding to vanishing and diverging *G* (see Labonte (2023) for further discussion).

**Figure 8:**
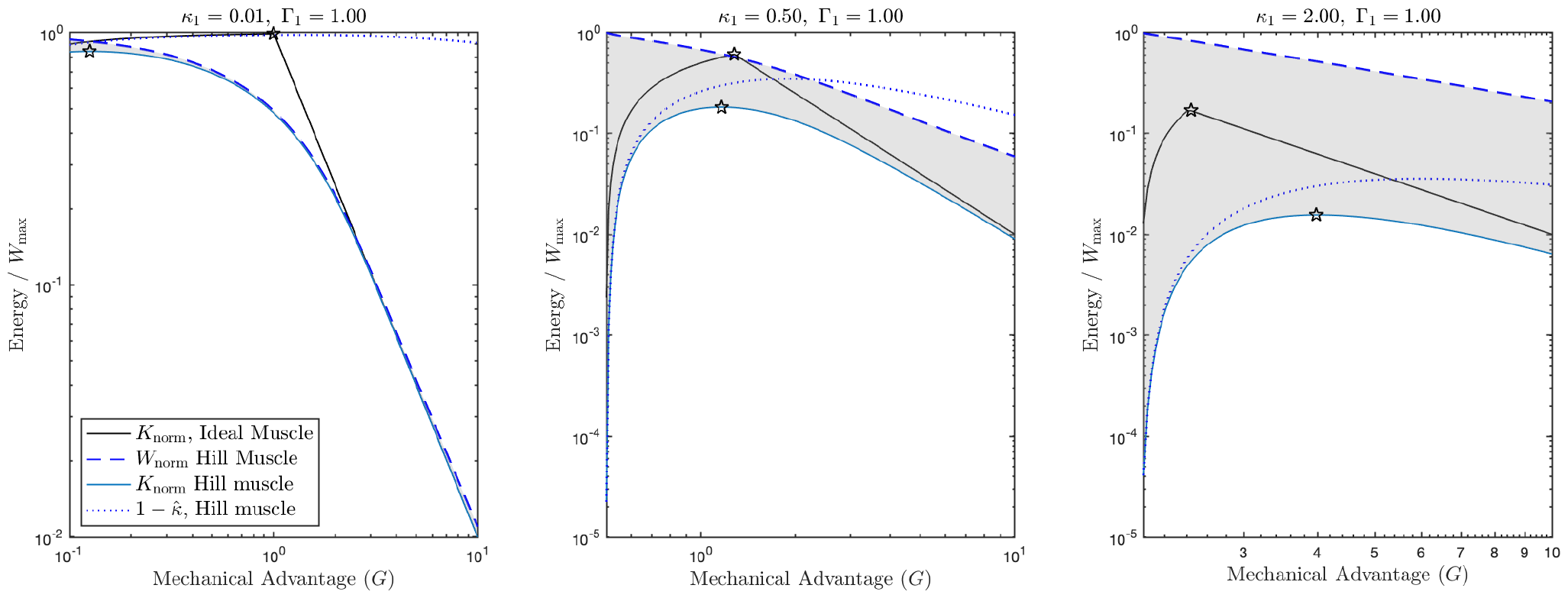
The energy landscapes for a Hill-muscle with a linear force-velocity relationship are qualitatively similar to those for an idealised muscle; the examples shown here are for contractions against a constant external force. From left to right, the ratio of external to muscle force *κ*_1_ increases; for all cases, Γ_1_ = 1. For both a Hill- and an idealised muscle, the work output (dashed line) decreases as *G* is increased. Reducing *G* by too much, in turn, reduces the fraction of work that flows into kinetic energy, the transmission efficiency (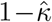, dotted line), and increases the time required to deliver each unit of work (not shown). Thus, although the optimal mechanical advantage will now have a different magnitude, it will still be intermediate. This result arises, because the energy outputs of Hill-muscle (blue solid line) and the idealised muscle (black solid line) are indistinguishable in the limits of vainishing and diverging *G*, respectively.

### The allometry of the mechanical advantage

The “effective mechanical advantage” that characterises the limbs of terrestrial mammals was extracted for 16 quadruped and 24 macropod species from Basu and Hutchinson (2022); Bennett and Taylor (1995); Biewener (1989b, 2005); Ren et al. (2010), as described in Labonte et al. (2024). Data on the mechanical advantage of the basalar muscle of the dragonfly flight apparatus was extracted from Schilder and Marden (2004), using Ankit Rohatgi’s WebPlotDigitizer (Rohatgi 2021). This paper reports in-lever, out-lever and basalar muscle mass for 47 individuals from eight species, but does not contain an explicit relationship between body mass and mechanical advantage. In order to obtain this relationship, basalar muscle mass and body mass were related via ordinary least square (OLS) regression of log10-transformed data. Mean estimates for the basalar muscle mass *m*_*m*_ for each species were extracted from Fig. 4, and the mean body mass *m*_*b*_ was provided in Tab. 1. OLS regression then yielded the relation *m*_*b*_ = 71*m*_*m*_. In-lever and out-lever were extracted from Fig. 5A and B, and lever pairs formed as accurately as possible, using the magnitude of the basalar muscle mass and species identification as guides.

**Table 1:**
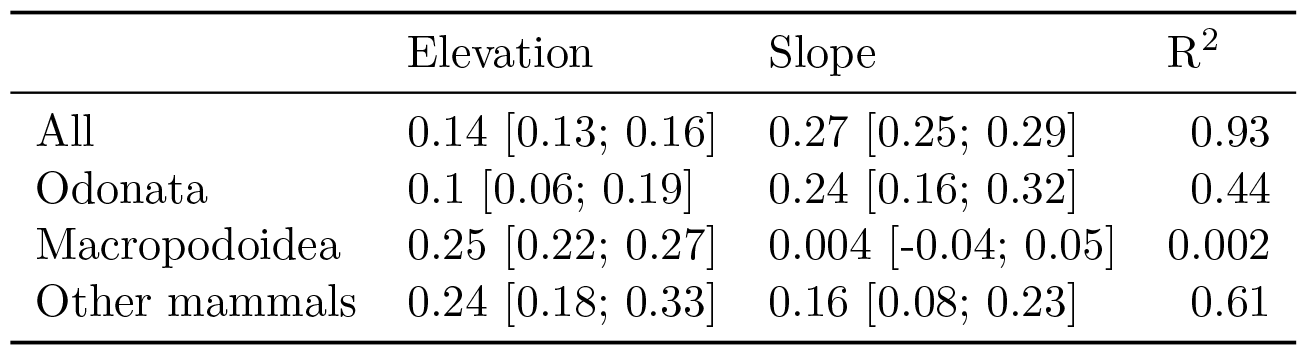
Results of ordinary least squares regression on log10-transformed data, with body mass in kilograms. Values in parentheses indicate 95% confidence intervals.

The relationship between body mass and mechanical advantage is influenced not only by mechanical and ecological constraints, but also by evolutionary history (Biewener 2005). In lieu of a phylogenetic regression, we roughly estimate the strength of the phylogenetic constraints by conducting four separate OLS regressions on log10-transformed data, summarised in Tab. 1: one including all data, one only for dragonflies, one only for macropods, and one for all mammalian quadrupeds. In order to directly estimate the optimal mechanical advantage for terrestrial mammals, we estimate Γ_1_ *≈* 0.0063mass^2*/*3^kg^*−*2*/*3^ and 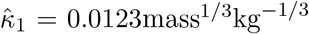 for the gravitational force, using the data provided in in Tab. 1 in Labonte et al. (2024). Eq. 15 then yields *G*_opt_ *≈* 1*/*11mass^1*/*3^.

Balance of aerodynamic forces yields a prediction 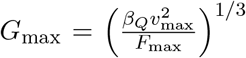, where *β*_*Q*_ is a quadratic drag multiplier (proportional to area). Using *β*_*Q*_ *∝m*^2*/*3^, *F*_max_ *∝m*^2*/*3^ and *v*_max_ *∝m*^1*/*3^, predicts *G*_max_ *∝*_*m*_^2*/*9^

### The magnitude of the optimal mechanical advantage

If muscle provides the dominant force in the system, the contraction is approximately inertial, and the magnitude of the optimal mechanical advantage follows as:

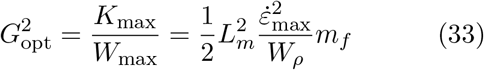

where *W*_*ρ*_ *≈* 70 J kg^-1^ is the work density of muscle, and *m*_*f*_ = *m*_*m*_*m*^*−*1^ is the ratio between muscle mass and payload mass (Labonte et al. 2024). For a representative maximum strain rate 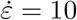 lengths per second, and a plausible value of *m*_*f*_ = 0.1, one may find:

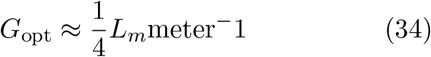

which is the estimate used in the main manuscript.

### Case studies

#### Praying mantis strike

The predatory strike of the raptorial forelimb of a praying mantis (*Heirodula membranacea*) provides an example of a rapid, approximately inertial movement. To determine the optimal mechanical advantage, only the ungeared physiological similarity index Γ_1_ needs be estimated.

Gray and Mill (1983) point to the Coxal trochanteral extensor and thoracic trochanteral extensor as the primary extensors of the coxotrochanteral joint. These muscle have a combined mass of 29.6 mg, yielding a work capacity of *W*_max_ = 2.1 mJ assuming a work density of 70 J/kg (Labonte et al. 2024). The trochanteral extensor has a characteristic fascicle length of *l*_*m*_ = 1.04 cm, and we assumed a maximum contraction speed of 10 lengths per second (Labonte et al. 2024).

To determine the moment of inertia of the forelimb distal to the coxa, the femur and trochanter were modeled as an ellipsoid, while the tibia was modeled as a cylinder, each with constant mass density of 1 g/cm^3^. Length dimensions were extracted from figure 6 in Gray and Mill (1983), with the same pose assumed, and out-of-plane thickness of the femur was taken as 1/9 femur+trochanter length. We neglect the inertia of the tarsus. This yielded a combined mass of 0.17 g, and radius of gyration of 1.85 cm.

Combination of these estimates results in Γ_1_ = 4.4 *×* 10^*−*4^, and thus 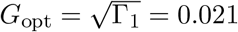. Gray and Mill (1983) report coxo-trochanteral moment arms between 0.04 to 0.1 cm during the strike; divided by the radius of gyration, this corresponds to *G*_emp_ *∈* [0.02, 0.05].

#### Kangaroo rat jump

The jump of a kangaroo rat serves as an example for a contraction against non-negligble gravitational forces. Rankin et al. (2018) present a musculoskeletal model of *Dipodomys deserti* from which relevant parameters can be obtained. The animal has a body mass 106 g and three ankle extensors per leg (Plantaris and lateral and medial Gastrocnemius) with combined mass, work capacity and maximum force of 2.63 g, 0.184 J and 68.8 N (summed across two legs). With a maximum strain of 0.3 (Labonte et al. 2024), this yields an effective fiber length of 18 mm. Javidi et al. (2020) report a maximum contraction velocity of 12 fiber lengths per second, which then yields Γ_1_ = 0.013. Assuming that the dominant external force is body weight, and using a gravitational acceleration of 9.81 m s^-2^, yields 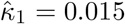.

Rankin et al. (2018) report maximum ankle extension moments of 146 to 167 N mm at 90 ankle flexion. This corresponds to in-lever moment arms of 4.24 to 4.85 mm, close to the values of 5.6 mm reported by Biewener and Blickhan (1988). For the outlever length, we assume the ground reaction force vector to be vertical with a center of pressure somewhere along the phalanges. Using model data presented by Rankin et al. (2018), the maximum outlever length was defined as (midfoot + toes) - (min in-lever), and the minimum outlever was defined as (midfoot - max in-lever). With a midfoot length of 24 mm and a toe length of 17.9 mm, this yields an empirical estimate of *G*_emp_ *∈* [0.11, 0.24].

#### Frog swimming

As an example for contractions against a drag force, we study the plantaris-driven ankle extension of *Xenopus laevis*, following Richards (2008); Richards and Sawicki (2012). From Richards and Sawicki (2012), we extract a frog of body mass of 30 g and a foot mass of 3 g. With this body mass, we use regression equations from Clemente and Richards (2013) to determine muscle length (*l*_*m*_ = 2.1 cm), foot area (*A* = 5.7 cm^2^), foot length (2.8 cm), maximum isometric force (*F*_max_ = 7.0 N), and maximum contraction velocity (*v*_max,*rel*_ = 6.8 lengths per second). The work capacity follows as the product between isometric force times muscle length with a maximum strain of 0.3 (Labonte et al. 2024), giving 0.0445 J. Using the trapezoidal approximation of foot area and length measurements from Richards (2008), and assuming a constant density, we calculate the moment of inertia of such a foot about the ankle as *I* = 0.01 kg cm^2^, and a center of area at *R* = 1.6 cm from the ankle. This gives a “mass equivalent” of *I/R*^2^ = 4 g. The ungeared physiological similarity index is therefore Γ_1_ = 8.7 *×* 10^*−*4^.

Following Richards and Clemente (2013b) we use a drag coefficient of *C*_*d*_ = 2 for the foot, and determine the quadratic drag multiplier *β*_*Q*_ = 1*/*2*ρAC*_*d*_ = 0.57 kg m^-1^, where *ρ* = 1000 kg m^-3^ is the density of water. The equivalent linear drag multiplier is thus *β*_*L*_ = 0.69 kg s^-1^ (Eq. 32). The maximum ungeared force ratio for linear drag follows as *κ*_max,1_= *β*_*L*_*l*_*m*_*v*_max,*rel*_*/F*_max_ ^= 0.014^.

The plantaris moment arm is also derived from a regression equation (Clemente and Richards 2013) as 0.16 cm. Compared to the center of area, this yields a mechanical advantage of 0.1. A lower mechanical advantage of 0.06 is reported by Richards and Clemente (2013b) in *Xenopus laevis*. These two values are taken as a sensible empirical range. We thus estimate *G*_emp_ *∈* [0.06, 0.1]

