## Supplemental Information for "Optimising the flow of mechanical energy in musculoskeletal systems through gearing"

Scholz MN, Bobbert MF, Knoek van Soest AJ. 2006. Scaling and jumping: Gravity loses grip on small jumpers. Journal of Theoretical Biology 240:554–561. doi:10.1016/j.jtbi.2005.10.015.

Simone Y, van Der Meijden A. 2018. Fast and fine versus strong and stout: a trade-off between chela closing force and speed across nine scorpion species. Biol J Linn Soc 123:208–217.

Smith JM, Savage RJG. 1956. Some locomotory adaptations in mammals. Zoological Journal of the Linnean Society 42:603–622.

Strobbe F, McPeek MA, De Block M, De Meester L, Stoks R. 2009. Survival selection on escape performance and its underlying phenotypic traits: a case of many-to-one mapping. Journal of Evolutionary Biology 22:1172–1182.

Thornton LH, Dick TJ, Hutchinson JR, Lichtwark GA, McGowan CP, Rubenson J, Wiktorowicz-Conroy A, Clemente CJ. 2024. Unlocking the secrets of kangaroo locomotor energetics: Postural adaptations underpin increased tendon stress in hopping kangaroos. bioRxiv : the preprint server for biology doi:10.1101/2024.02.05.578950.

Tseng ZJ, Garcia-Lara S, Flynn JJ, Holmes E, Rowe TB, Dickson BV. 2023. A switch in jaw form-function coupling during the evolution of mammals. Philosophical Transactions of the Royal Society B: Biological Sciences 378:20220091. doi: 10.1098/rstb.2022.0091.

Usherwood JR. 2013. Constraints on muscle performance provide a novel explanation for the scaling of posture in terrestrial animals. Biology Letters 9:20130414. doi:10.1098/rsbl.2013.0414.

Wainwright PC. 2007. Functional versus morphological diversity in macroevolution. Annu. Rev. Ecol. Evol. Syst. 38:381–401.

Wainwright PC, Alfaro ME, Bolnick DI, Hulsey CD. 2005. Many-to-one mapping of form to function: A general principle in organismal design?1. Integr Comp Biol 45:256–262. doi:10.1093/icb/45.2.256.

Westneat MW. 1994. Transmission of force and velocity in the feeding mechanisms of labrid fishes (teleostei, perciformes). Zoomorphology 114:103–118.

Westneat MW. 2004. Evolution of levers and linkages in the feeding mechanisms of fishes1. Integr Comp Biol 44:378–389.

Wilson AM, Hubel TY, Wilshin SD, Lowe JC, Lorenc M, Dewhirst OP, Bartlam-Brooks HLA, Diack R, Bennitt E, Golabek KA, Woledge RC, McNutt JW, Curtin NA, West TG. 2018. Biomechanics of predator-prey arms race in lion, zebra, cheetah and impala. Nature 554:183–188.

Wilson JW, Mills MGL, Wilson RP, Peters G, Mills MEJ, Speakman JR, Durant SM, Bennett NC, Marks NJ, Scantlebury M. 2013. Cheetahs, acinonyx jubatus, balance turn capacity with pace when chasing prey. Biology Letters 9:20130620. doi:10.1098/rsbl.2013.0620.

### Supplementary Information

#### Optimal gearing for velocity-dependent external forces

#### General procedure

The external force $F_e$ may depend on the velocity, $F_e(u)$. A common biological example of velocity-dependent external force is the drag force, $F_D$. Drag is a complex physical phenomenon, and a first-order model typically considers a drag force that varies either linearly or quadratically with speed. A linear drag force, $F_D \propto u$, is an appropriate model for viscous drag, which dominates at small Reynolds numbers (that is, slow speed, high viscosity, or small length). At intermediate to large Reynolds numbers, a quadratic drag force, $F_D \propto u^2$, is more appropriate.

The general analysis for both a linear and quadratic drag force proceeds from the conservation of energy:

$$\begin{aligned} W =& \Delta K \\ \int (F_o - F_D)\, dx =& m \int u du \end{aligned} \tag{22}$$

Because the drag force depends on $u$, solution requires separation of variables:

$$\int F_o dx = m \int \frac{u}{1 - F_D/F_o} du \tag{23}$$

To introduce the mechanical advantage, we use the couplings $F_0 = F_i G$, and $x = \delta/G$, and note that $G$ is optimal if the maximum displacement and shortening speed of muscle are reached simultaneously:

$$\int^{\delta_{\max}} F_i d\delta = m \int^{u_{max}} \frac{u}{1 - F_D/(F_i G_{\text{opt}})} du \tag{24}$$

For a linear drag force, $F_{D,L} = \beta_L u$, the velocity integral evaluates to:

$$\begin{aligned} W_{\max} =& m \int_0^{u_{\max}} \frac{u}{1 - u\, \beta_L/(F_i G_{\text{opt}})} du \\ \frac{\kappa_{\max,1}}{\Gamma_1} =& -2 \left[ 1 + \frac{G_{\text{opt}}^2}{\kappa_{\max,1}} \log \left( 1 - \frac{\kappa_{\max,1}}{G_{opt}^2} \right) \right] \end{aligned} \quad (25)$$

where we used the coupling $u = v/G$, and introduced $\kappa_{\max,1} = \beta_L v_{\max} F_{\max}^{-1}$ as the ratio of the maximum drag and driving force for an ungeared muscle with a mechanical advantage of unity.

For a quadratic drag force, $F_{D,Q} = \beta_Q u^2$, one finds instead:

$$\begin{aligned} W_{\max} =& m \int_0^{u_{\max}} \frac{u}{1 - u^2\, \beta_Q/(F_i G_{\text{opt}})} du \\ \frac{\kappa_{\max,1}}{\Gamma_1} =& -G_{\text{opt}} \log \left( 1 - \frac{\kappa_{\max,1}}{G_{opt}^3} \right) \end{aligned} \quad (26)$$

where $\kappa_{\max,1} = \beta_Q v_{\max}^2 F_{\max}^{-1}$ has the same physical meaning as before.

Thus, for both a linear and a quadratic drag force, the optimal mechanical advantage is determined uniquely by two dimensionless numbers, $\Gamma_1$ and $\kappa_{\max,1}$.

##### The optimal mechanical advantage for linear drag

For the linear drag force, an explicit solution for $G_{opt}$ can be found:

$$G_{\text{opt}} = \frac{\sqrt{\kappa_{\max,1}} \sqrt{\frac{\kappa_{\max,1}}{2\Gamma_1} + 1}}{\sqrt{\frac{\kappa_{\max,1}}{2\Gamma_1} + 1 + \mathrm{W}\left[ -\exp\left( -\frac{\kappa_{\max,1}}{2\Gamma_1} - 1 \right) \left( 1 + \frac{\kappa_{\max,1}}{2\Gamma_1} \right) \right]}} \quad (27)$$

where $W$ is the Lambert W function. It is hard to develop an intuitive feel for this expression, but the limits provide a clear physical picture. For convenience, we define $\gamma = \kappa_{\max,1}/(2\Gamma_1)$, and consider the limits $\gamma \to 0$ (inertial forces dominate) and $\gamma \to \infty$ (external force dominate). Through Taylor expansion about $\gamma = 0$, we find $\lim_{\gamma \to 0} 1 + \mathrm{W}\left[ -\exp\left( -\gamma - 1 \right) \left( 1 + \gamma \right) \right] = \gamma$. The right-side limit for the remaining term is:

$$\lim_{\gamma \to 0} G_{\text{opt}} = \sqrt{\Gamma_1} \quad (28)$$

In this limit, inertial forces dominate, and optimal mechanical advantage is thus independent of external forces. For $\gamma \to \infty$, the productlog-term goes to 0, which yields:

$$\lim_{\gamma \to \infty} G_{\text{opt}} = \sqrt{\kappa_{\max,1}} \quad (29)$$

The inertial force is now irrelevant, and the optimal mechanical advantage is the value of $G$ which ensures an equilibrium of the parasitic and driving force.

For any intermediate value, the optimal gear ratio depends on a complex combination of the three key forces: the inertial force, the driving force, and the parasitic force.

##### The optimal mechanical advantage for quadratic drag

To the best of our judgement, the implicit expression for $G_{\text{opt}}$ for a quadratic drag force allows no explicit writing. Instead, $G_{\text{opt}}$ has to be determined numerically. However, the limits can still be assessed, and follow in direct analogy to the case for a linear drag force.

When the drag force is large, $\gamma = \kappa_{\max,1}/(\Gamma_1) \to \infty$, and eq. 26 yields:

$$\lim_{\gamma \to \infty} G_{\text{opt}} = \left( \frac{\beta_Q v_{\max}^2}{F_{\max}} \right)^{1/3} = \sqrt[3]{\kappa_{\text{Qmax},1}} \quad (30)$$

which is equivalent to equilibrium of maximum dynamic forces (see also Richards and Clemente (2013b)):

$$\begin{aligned} F_{\max,o} =& F_{\max,D} \\ G_{\text{opt}} F_{\max} =& \beta_Q u_{\max}^2 \\ G_{\text{opt}} =& \left( \frac{\beta_Q v_{\max}^2}{F_{\max}} \right)^{1/3} \end{aligned} \quad (31)$$

When inertial forces dominate, $G_{\text{opt}} \to \sqrt{\Gamma_1}$, which follows as before.

Recognizing that both quadratic and linear drag have the same inertial limit for $G_{\text{opt}}$, the symbolic results for linear drag above can be used to approximate quadratic drag by setting the upper $G_{\text{opt}}$ limits to be equal. This implies:

The relationship between body mass and mechanical advantage is influenced not only by mechanical and ecological constraints, but also by evolutionary history (Biewener 2005). In lieu of a phylogenetic regression, we roughly estimate the strength of the phylogenetic constraints by conducting four separate OLS regressions on log10-transformed data, summarised in Tab. 1: one including all data, one only for dragonflies, one only for macropods, and one for all mammalian quadrupeds. In order to directly estimate the optimal mechanical advantage for terrestrial mammals, we estimate $\Gamma_1 \approx 0.0063\text{mass}^{2/3}\text{kg}^{-2/3}$ and $\hat{\kappa}_1 = 0.0123\text{mass}^{1/3}\text{kg}^{-1/3}$ for the gravitational force, using the data provided in in Tab. 1 in Labonte et al. (2024). Eq. 15 then yields $G_{\text{opt}} \approx 1/11\text{mass}^{1/3}$.

Balance of aerodynamic forces yields a prediction $G_{\max} = \left(\frac{\beta_Q v_{\max}^2}{F_{\max}}\right)^{1/3}$, where $\beta_Q$ is a quadratic drag multiplier (proportional to area). Using $\beta_Q \propto m^{2/3}$, $F_{\max} \propto m^{2/3}$ and $v_{\max} \propto m^{1/3}$, predicts $G_{\max} \propto m^{2/9}$

#### The magnitude of the optimal mechanical advantage

If muscle provides the dominant force in the system, the contraction is approximately inertial, and the magnitude of the optimal mechanical advantage follows as:

$$G_{\text{opt}}^2 = \frac{K_{\max}}{W_{\max}} = \frac{1}{2} L_m^2 \frac{\dot{\varepsilon}_{\max}^2}{W_\rho} m_f \tag{33}$$

where $W_\rho \approx 70\,\text{J kg}^{-1}$ is the work density of muscle, and $m_f = m_m m^{-1}$ is the ratio between muscle mass and payload mass (Labonte et al. 2024). For a representative maximum strain rate $\dot{\varepsilon} = 10$ lengths per second, and a plausible value of $m_f = 0.1$, one may find:

$$G_{\text{opt}} \approx \frac{1}{4} L_m \text{meter}^{-}1 \tag{34}$$

which is the estimate used in the main manuscript.

#### Case studies

##### Praying mantis strike

The predatory strike of the raptorial forelimb of a praying mantis (*Heirodula membranacea*) provides an example of a rapid, approximately inertial movement. To determine the optimal mechanical advantage, only the ungeared physiological similarity index $\Gamma_1$ needs be estimated.

Gray and Mill (1983) point to the Coxal trochanteral extensor and thoracic trochanteral extensor as the primary extensors of the coxo-trochanteral joint. These muscle have a combined mass of 29.6 mg, yielding a work capacity of $W_{\max} = 2.1$ mJ assuming a work density of 70 J/kg (Labonte et al. 2024). The trochanteral extensor has a characteristic fascicle length of $l_m = 1.04$ cm, and we assumed a maximum contraction speed of 10 lengths per second (Labonte et al. 2024).

To determine the moment of inertia of the forelimb distal to the coxa, the femur and trochanter were modeled as an ellipsoid, while the tibia was modeled as a cylinder, each with constant mass density of

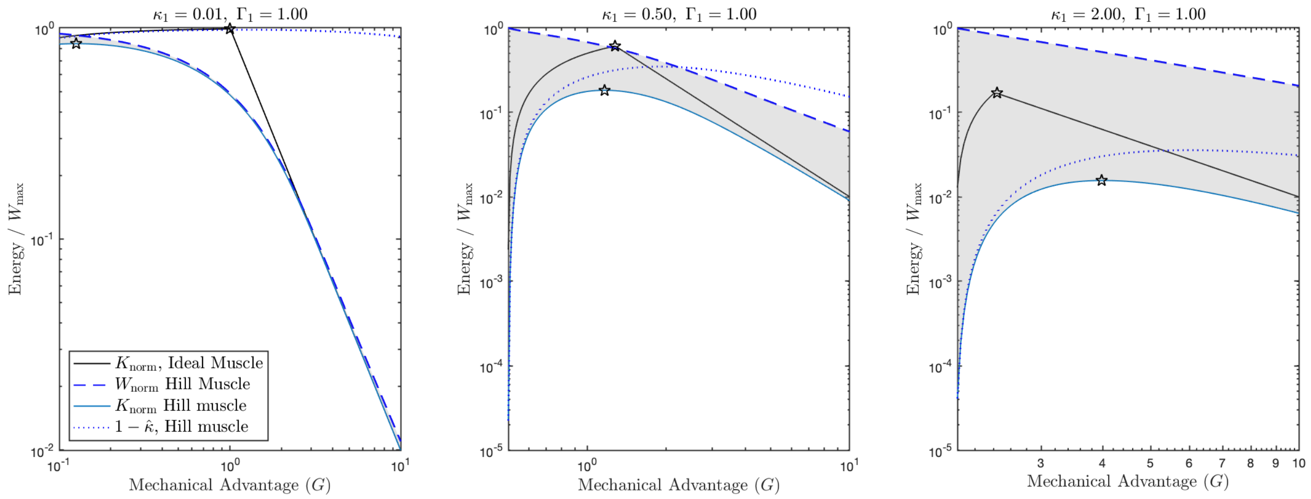


Figure 8: The energy landscapes for a Hill-muscle with a linear force-velocity relationship are qualitatively similar to those for an idealised muscle; the examples shown here are for contractions against a constant external force. From left to right, the ratio of external to muscle force $\kappa_1$ increases; for all cases, $\Gamma_1 = 1$. For both a Hill- and an idealised muscle, the work output (dashed line) decreases as $G$ is increased. Reducing $G$ by too much, in turn, reduces the fraction of work that flows into kinetic energy, the transmission efficiency ($1-\hat{\kappa}$, dotted line), and increases the time required to deliver each unit of work (not shown). Thus, although the optimal mechanical advantage will now have a different magnitude, it will still be intermediate. This result arises, because the energy outputs of Hill-muscle (blue solid line) and the idealised muscle (black solid line) are indistinguishable in the limits of vainishing and diverging $G$, respectively.

Table 1: Results of ordinary least squares regression on log10-transformed data, with body mass in kilograms. Values in parentheses indicate 95% confidence intervals.

| | Elevation | Slope | $R^2$ |
|---|---|---|---|
| All | 0.14 [0.13; 0.16] | 0.27 [0.25; 0.29] | 0.93 |
| Odonata | 0.1 [0.06; 0.19] | 0.24 [0.16; 0.32] | 0.44 |
| Macropodoidea | 0.25 [0.22; 0.27] | 0.004 [-0.04; 0.05] | 0.002 |
| Other mammals | 0.24 [0.18; 0.33] | 0.16 [0.08; 0.23] | 0.61 |

1 g/cm$^3$. Length dimensions were extracted from figure 6 in Gray and Mill (1983), with the same pose assumed, and out-of-plane thickness of the femur was taken as 1/9 femur+trochanter length. We neglect the inertia of the tarsus. This yielded a combined mass of 0.17 g, and radius of gyration of 1.85 cm.

Combination of these estimates results in $\Gamma_1 = 4.4 \times 10^{-4}$, and thus $G_{\text{opt}} = \sqrt{\Gamma_1} = 0.021$. Gray and Mill (1983) report coxo-trochanteral moment arms between 0.04 to 0.1 cm during the strike; divided by the radius of gyration, this corresponds to $G_{\text{emp}} \in [0.02, 0.05]$.

Following Richards and Clemente (2013b) we use a drag coefficient of $C_d = 2$ for the foot, and determine the quadratic drag multiplier $\beta_Q = 1/2\rho A C_d = 0.57$ kg m$^{-1}$, where $\rho = 1000$ kg m$^{-3}$ is the density of water. The equivalent linear drag multiplier is thus $\beta_L = 0.69$ kg s$^{-1}$ (Eq. 32). The maximum ungeared force ratio for linear drag follows as $\kappa_{\max,1} = \beta_L l_m v_{\max,rel}/F_{\max} = 0.014$.

The plantaris moment arm is also derived from a regression equation (Clemente and Richards 2013) as 0.16 cm. Compared to the center of area, this yields a mechanical advantage of 0.1. A lower mechanical advantage of 0.06 is reported by Richards and Clemente (2013b) in *Xenopus laevis.* These two values are taken as a sensible empirical range. We thus estimate $G_{\mathrm{emp}} \in [0.06, 0.1]$
